# The Aryl Hydrocarbon Receptor Controls IFNγ-Induced Immune Checkpoints PD-L1 and IDO via the JAK/STAT Pathway in Lung Adenocarcinoma

**DOI:** 10.1101/2024.08.12.607602

**Authors:** Megan Snyder, Zhongyan Wang, Brian Lara, Jocelyn Fimbres, Tachira Pichardo, Sarah Mazzilli, Mohammed Muzamil Khan, Vinay K. Duggineni, Stefano Monti, David H. Sherr

## Abstract

While immunotherapy has shown efficacy in lung adenocarcinoma (LUAD) patients, many respond only partially or not at all. One limitation in improving outcomes is the lack of a complete understanding of immune checkpoint regulation. Here, we investigated a possible link between an environmental chemical receptor implicated in lung cancer and immune regulation, (the aryl hydrocarbon receptor/AhR), a known but counterintuitive mediator of immunosuppression (IFNγ), and regulation of two immune checkpoints (PD-L1 and IDO). AhR gene-edited LUAD cell lines, a syngeneic LUAD mouse model, bulk- and scRNA sequencing of LUADs and tumor-infiltrating leukocytes were used to map out a signaling pathway leading from IFNγ through the AhR to JAK/STAT, PD-L1, IDO, and tumor-mediated immunosuppression. The data demonstrate that: **1)** IFNγ activation of the JAK/STAT pathway leading to PD-L1 and IDO1 upregulation is mediated by the AhR in murine and human LUAD cells, **2)** AhR-driven IDO1 induction results in the production of Kynurenine (Kyn), an AhR ligand, which likely mediates an AhR➔IDO1➔Kyn➔AhR amplification loop, **3)** transplantation of AhR-knockout LUAD cells results in long-term tumor immunity in most recipients. **4)** The 23% of AhR-knockout tumors that do grow do so at a much slower pace than controls and exhibit higher densities of CD8^+^ T cells expressing markers of immunocompetence, increased activity, and increased cell-cell communication. The data definitively link the AhR to IFNγ-induced JAK/STAT pathway and immune checkpoint-mediated immunosuppression and support the targeting of the AhR in the context of LUAD.

## Introduction

Lung cancer is one of the most commonly diagnosed and deadliest of human malignancies (1). More than half of all lung cancers are adenocarcinomas (LUAD), a subset of non-small cell lung cancers (NSCLCs) likely derived from alveolar type 2 cells (2). Smoking is the single greatest LUAD risk factor. However, LUAD is also the most common form of lung cancer seen in never-smokers (3). Surgery, radiotherapy, and chemotherapy remain front-line treatments for LUAD, with varying degrees of success. More recently, immunotherapy, particularly with monoclonal antibodies targeting the PD-1/PD-L1 axis either as monotherapy or in combination with traditional treatments, has proven effective in subsets of LUAD patients (4, 5). However, the proportion of patients eligible for immunotherapy remains limited and many patients respond only partially or not at all. Importantly, not all factors controlling expression of the immune checkpoints targeted by immunotherapy have been defined. That said, it has been shown that IFNγ, usually associated with positive immune responses, can contribute to immune suppression by upregulating PD-L1 and the indoleamine-2,3-dioxygenases, IDO1 and IDO2, proximal and redundant rate-limiting enzymes in the kynurenine (Kyn) pathway of tryptophan metabolism (6). Kyn itself, produced by IDO^+^ melanomas (7), ovarian (8) (9), squamous cell (10), and colon (11) carcinomas induces potent immunosuppression in the tumor microenvironment (TME). At least one pathway through which IFNγ induces these immune checkpoints is the JAK/STAT pathway (12, 13). Therefore, it is important to identify factors that regulate the IFN-activated JAK/STAT pathway and lead to immune checkpoint expression and tumor-mediated immunosuppression. As shown herein, one such factor is the AhR.

The AhR is a ligand-activated transcription factor and protein-binding partner originally recognized for its activation by environmental chemicals including 2,3,7,8-tetrachlorodibenzo-p-dioxin (TCDD), polychlorinated biphenyls (PCBs), and planar polycyclic aromatic hydrocarbons (PAH). AhR activation induces expression of CYP1A1, CYP1A2, and CYP1B1 monoxygenases capable of metabolizing some environmental AhR ligands, including PAH common in cigarette smoke, into mutagenic intermediates (14). These smoke-derived mutagens have long been associated with LUAD and other cancers (15, 16). Furthermore, and more germane to the present studies, many of these same environmental AhR ligands are highly immunosuppressive (17, 18) and the AhR itself, however it is activated, is associated with immunosuppression in several contexts (19–23).

The effects of environmental chemicals aside, accumulating evidence implicates the AhR in cancer even in the absence of environmental ligands (6, 24–26). Thus, the AhR is hyper-expressed and chronically active in several cancers (27). It is now apparent that endogenous AhR agonists are at least partially responsible for this activity and that they drive malignant cell migration, metastasis, and cancer stem cell properties (28–31). Indeed, the level of AhR activity is inversely related to survival in lung cancer patients (32). While AhR ligands may derive from multiple sources, including the diet and microbiome (33, 34), some endogenous ligands originate from within the TME itself (6, 27). One source of such agonists is the IDO-dependent Kyn pathway of tryptophan metabolism. Notably, the AhR can upregulate expression of IDO1/2 which generates AhR ligands, including Kyn itself, through the Kyn metabolic pathway. Kyn-activated AhR also has been implicated in PD-1 expression on tumor infiltrating T cells (7), and on PD-L1 expression on primary human lung epithelial cells (35) and oral squamous cell carcinomas (36).

The apparent influence of IFNγ and the AhR on IDO levels, Kyn production, and immune checkpoint expression suggests the existence of an intricate pathway of interactions involving IFNγ, AhR, IDO, Kyn (or its downstream metabolites/AhR ligands), and PD-L1 resulting in suppression of tumor immunity in the TME. Here, these interactions were more clearly mapped using *AhR* knockout murine (CMT167) human (A549) LUAD cells, a syngeneic LUAD mouse model, immunophenotyping, and bulk and single cell RNA sequencing of whole LUAD and sorted tumor-infiltrating leukocytes respectively. Surprisingly, the results indicate a novel pathway within LUAD cells in which the AhR controls IFN type II-induced JAK/STAT signaling leading to IDO1/2 and PD-L1/PD-L2 expression and, ultimately, immunosuppression in the TME. Collectively, the results reveal the AhR to be a master regulator of IFNγ signaling and help explain mechanisms of immune checkpoint regulation, including the counterintuitive role that IFN plays in immunosuppression in the LUAD context.

## Materials and Methods

### Cell lines and Cell Culture

Murine CMT167 lung adenocarcinoma cell line (“CMT167”) were kindly provided by Dr. Raphael Nemenoff (University of Colorado-Denver). CMT167 cells were selected since they have commonly been used to study immune checkpoints in LUAD and since they harbor a KrasG12V mutation, one of the most common driver mutations in human NSCLC (37). Human A549 LUAD cells were obtained from the American Type Culture Collection (ATCC). The A549 cell line was chosen in part because it harbors a KrasG12S mutation, another common driver mutation in human LUADs, and because it has been used extensively to study regulation of immune checkpoints (38, 39). Cells were cultured in Dulbecco’s Modified Eagle Medium (Corning Inc., Corning, NY) supplemented with 10% fetal bovine serum (Gemini Bioproducts LLC, West Sacramento, CA), 1% penicillin/streptomycin (Life Technologies, Gaithersburg, MD), and 1% L-glutamine (Fisher Scientific, Hampton, NH) at 37°C with 5% carbon dioxide. Cells were kept in culture no longer than eight weeks and new aliquots were thawed periodically. Cultures were confirmed to be mycoplasma negative every two months.

For *in vitro* RT-qPCR, western immunoblotting, and immunophenotyping experiments CMT167 or A549 cell lines (50,000 cells/well) were cultured in 6-or 12-well plates for 24-72 hours with or without 1-100 ng/ml IFNγ (PeproTech, Cranbury, NJ), 10 µM benzo(a)pyrene (B(a)P)(Sigma-Aldrich, Burlington, MA), or 0.5 µM 6-formylindolo(3,2-b)carbazole (FICZ)(Sigma-Aldrich, Burlington, MA). Each experiment included a minimum of three replicates, with each replicate representing a pool of at least two wells.

### CRISPR/Cas9-mediated knockouts

Single-guide RNAs (sgRNAs) targeting the mouse *AhR* gene (Exon1), the mouse *Ifnγr1* gene (Exon1) and the mouse *Cd274 gene* (Exon1) were designed using the web resource (https://www.synthego.com/products/bioinformatics/crispr-design-tool). Two individual sgRNAs were used to target the *AhR* gene (sgRNA1, 5′-CGGCTTGCGCCGCTTGCGGC −3′; sgRNA2, 5′-AAACGTGAGTGACGGCGGGC-3′).. Complementary oligonucleotides for sgRNAs were annealed, and cloned into the sgOpti vector (Addgene, Cambridge, MA, #85681) *BsmBI* sites (40) using standard procedures (40). Lentivirus particles were generated in the HEK293NT cells by co-transfecting the lentiCas9-Blast (Addgene, #52962), AhR-sgOpti, Ifnγr1-sgOpti or Cd274-sgOpti plasmids, and the packaging plasmids (pLenti-P2A and pLenti-P2B, Cat. # LV003, Applied Biological Materials Inc. Richmond, BC, Canada), using lipofectamine 2000 (Invitrogen) according to the manufacturer’s instructions. Lentiviral particle-containing supernatants were collected 24 and 48h after transfection and filtered using a 0.45-μm filter. CMT167 cells were transduced with the lentivirus in the presence of 5 µg/ml polybrene. Forty-eight hours after transduction, cells were selected with Blasticidin (5 µg/ml) and Puromycin (4 µg/ml) for two weeks. Gene deletion was validated by DNA sequencing (not shown), western immunoblotting (**Supplemental Fig. 1A,D**) and by lack of transcriptional responsiveness (i.e., *CYP1B1* induction) in response to a strong AhR ligand (**Supplemental Fig. 1B,E**). Two AhR knockout CMT167 clones, C1 and D2 (CMT167^AhR-KO^), and two AhR knockout A549 clones, P15 and P16 (A549^AhR-KO^), were identified. Control lines (CMT167^Cas9^) were generated with the Cas9 vector without guide RNA.

### Colorimetric kynurenine assay

Culture supernatants (160 μl) were transferred to 96-well round-bottom plates and mixed with 10 µl / well 30% (v/v) freshly prepared trichloroacetic acid. The plates were incubated at 50°C for 30 minutes to hydrolyze N-formyl-kynurenine to kynurenine. Samples were centrifuged at 3,000 g for 10 min. Aliquots of the supernatant (100 ml) were transferred to flat-bottom 96-well plates and mixed with 100 µl freshly prepared Ehrlich’s reagent (1.2% w/v 4-dimethylamino-benzaldehyde in glacial acetic acid). Absorbance at 492 nm was measured using a microplate reader (BioTek, Winooski, VT, USA) with the Gen5 software (BioTek) after a 10 min incubation. Kynurenine concentrations were calculated by reference to a standard kynurenine curve.

### RNA extraction and bulk RNA sequencing

CMT167^AhR-KO^, A549^AhR-KO^, wildtype (CMT167^WT^) or CMT^Cas9^ cells, in triplicate or quadruplicate wells, were harvested and total RNA was extracted using the RNeasy Plus Mini Kit (QIAGEN, Hilden, Germany) according to the manufacturer’s instructions. cDNA was generated using the High-Capacity cDNA Reverse Transcription Kit (Applied Biosystems, Waltham, MA) following the manufacturer’s instructions. cDNA was sequenced using the Boston University Microarray and Sequencing Core Facility. Between 18 and 28 million total read pairs (CMT167 lines) or 50 to 71 million total read pairs (A549 lines) were obtained per sample. Quality metrics were similar across all samples with no technical outliers. The Broad Institute’s Morpheus software (https://software.broadinstitute.org/morpheus, Broad Institute, Cambridge, MA) was used to generate heat maps. QIAGEN’s Ingenuity Pathway Analysis (IPA) software https://qiagenbioinformatics.com/products/ingenuity-pathway-analysis, QIAGEN, Inc.) was used to perform regulation pathway analyses.

### RT-qPCR

Reverse Transcription-quantitative polymerase chain reaction (RT-qPCR) analysis was conducted with the QuantStudio 3 Real-Time PCR System (Thermo Fisher Scientific, Waltham, MA). Relative mRNA expression was quantified using the comparative Ct (ΔΔCt) method according to the ABI manual (Applied Biosystems). Amplification of glyceraldehyde-3-phosphate dehydrogenase (*Gapdh*) served as an internal reference in each reaction. Assays were performed in triplicate. The following TaqMan assays were purchased from Thermo Fisher Scientific: *Cd274* (Mm03048248_m1), *Ido1* (Mm00492590_m1), *Ido2* (Mm00524210_m1), *Cyp 1a1* (Mm00487218_m1), *Cyp1b1* (Mm00487229_m1), *Jak2* (Mm01208489_m1), *Stat1* (Mm012 57286_m1), *gapdh* (Mm99999915_g1), *CD274* (Hs00204257_m1), *IDO1* (Hs00984148_m1), *J AK2* (Hs01078136_m1), *STAT1* Hs01013996_m1), *STAT3(*<u>Hs00374280_m1</u>), *Muc1* (<u>Mm00449604_m1</u>), *Col5a1* (<u>Mm00489299_m1</u>), *Thbs1* (<u>Mm01335418_m1</u>), *Egfr* (<u>Mm01187858_m1</u>), *Itgb2* (<u>Mm00434513_m1</u>), *Cd109* (<u>Mm00462151_m1</u>), *Ccl2* (<u>Mm00441242_m1</u>), *Ccl5* (<u>Mm01302427_m1</u>),*GAPDH* (Hs99999905_m1).

### Western blotting

Cells were grown to 70-80% confluence, harvested with trypsin, and lysed with radioimmunoprecipitation assay (RIPA) buffer with protease inhibitors. Blots were incubated overnight with AhR-(1:1000, #MA1-514, Thermo Fisher Scientific), PD-L1-(1:1000, ab213480, Abcam), or IDO1-(1:1000, #51851, Cell Signaling) specific antibodies and with β-actin-(1:2000, # A5441, Sigma-Aldrich) or GAPDH-(1:1000, #97166, Cell Signaling) specific antibody to serve as a loading control.

### *In vivo* experiments

Six to eight week-old C57BL/6J mice (Jackson Laboratories) were anesthetized and the right flank injected subcutaneously with CMT167^WT^ or CMT167^AhR-KO^ cells using a 1 ml syringe and 27½ gauge needle with 10^6^ cells in Hank’s Balanced Salt Solution. The *in vitro* growth rates of CMT167^AhR-KO^ clones C1 and D2 and A549^AhR-KO^ clones P15 and P16 and their respective WT and Cas9 controls were not significantly different. Average tumor size per mouse was determined using a caliper and tumor volume calculated as (width^2^ x length)/2. Tumors were excised by cutting the surrounding fascia with scissors to separate the largely round tumor tissue suspended between the dermis and peritoneum. For rechallenge experiments, 10^6^ CMT167^WT^ cells were injected into the left flank 65 days after a previous injection of CMT167^AhR-KO^ cells into the right flank. Euthanasia was performed when one dimension of the tumor reached 20 mm or when ulceration without fibrin crust formation was seen.

### Tumor infiltrating leukocyte (TIL) isolation

Single cell suspensions of tumor-infiltrating leukocytes were obtained by chopping and digesting tumors in 1.25 mg/mL collagenase IV (STEMCELL Technologies, Cambridge, MA), 0.025 mg/mL hyaluronidase (Sigma-Aldrich), and 0.01 mg/mL DNAse I (Sigma-Aldrich) followed by washing in HBSS and processing using the Miltenyi GentleMACS system (Miltenyi, Bergisch Gladbach, Germany). Digests were then filtered using 70 µm cell strainers. For some experiments, (e.g., single cell RNA sequencing, intracellular cytokine staining) digests were further cleaned by magnet bead separation of dead cells from live cells using Miltenyi’s Dead Cell Removal Kit per manufacturer’s instructions.

### Flow cytometry

Live cells in single cell suspensions were identified using a Fixable Live/Dead stain (Biolegend, San Diego, California) followed by staining according to the manufacturer’s protocol. For intracellular markers, cells were first fixed and permeabilized (eBioscience, Waltham, MA). All antibodies were purchased from Biolegend or Thermo Fisher Scientific except for the unconjugated anti-L-kynurenine antibody which was obtained from ImmuSmol (Bordeaux, France). All surface markers were stained at concentrations between 1:50 to 1:200, and all intracellular markers were stained at a concentration of 1:100 to 1:200. L-kynurenine staining required a secondary antibody which was used at a 1:100 concentration. Assays were analyzed using the Cytek Aurora flow cytometer (Cytek Biosciences, Freemont, CA). Representative CD4, CD8, CD44, and PD1 plots are provided in **Supplemental Figure 2**.

### Immunofluorescence

Subcutaneous lung adenocarcinomas and whole lungs were fixed in 10% neutral buffered formalin for 24-48h, processed and embedded in paraffin before being cut into 5 µm sections. Mounted slides were used immediately or stored at −80^0^ C to preserve antigens for immunofluorescence. Formalin fixed, paraffin embedded sections were deparaffinized in xylene baths followed by ethanol bath rehydration. Antigen retrieval was performed in pH 10 Tris-HCL buffer (Sigma-Aldrich) by microwaving samples for 10 min at 20% power. Samples were then blocked for 1 hour in 5% goat serum and PBS and then incubated overnight at 4° C with CD45-specific (Abcam, Cat# ab154885, 1:100) or AhR-specific (Abcam Cat# ab213480, 1:50) primary antibodies in blocking buffer. After washing in PBS + 0.1% Tween20 (PBST), AlexaFluor® 488-labelled secondary antibody (AbcamA27034, 1:400) was added and samples were incubated for 1 hour in blocking buffer. After washing with PBST, slides were sealed with DAPI Prolong Gold, dried and imaged within two days on a Zeiss Axioscan.Z1 Slide Scanner. Fluorescent quantification was done by selecting the whole tissue sections, detecting cells from the DAPI channel and loading antibody channel classifiers in QuPath.

### Single Cell RNA sequencing (scRNA-seq) and analysis

CMT167^WT^ or CMT167^AhR-KO^ tumors were digested as above and dead cells removed by magnetic bead separation using Miltenyi’s Dead Cell Removal Kit. Single cell suspensions were sorted for live CD45^+^ cells and resuspended in 0.04% BSA (Sigma). Viability was determined manually by trypan blue exclusion and cells were shown to be ≥90% viable. An average of 5,792 viable cells from wildtype tumors and an average of 3,173 viable cells from AhR-KO tumors were loaded into an Illumina cartridge in the Boston University Microarray and Single Cell Sequencing Core Facility. Barcoding and scRNA-seq cDNA library preparation were done using the Chromium platform from 10X Genomics in accordance with the manufacturer’s guide. Sequencing was done using the Illumina NexSeq2000 System. Over 150 million reads were obtained per sample.

Single cells were preprocessed using singlecellTK (41) by applying doublet detection (aggregation of four methods; scDblFinder, cxds, bcds, doubletFinder), decontamination (5%), and filtering for mitochondrial gene content (>20%). Low cell-gene content was filtered (<400 genes per cell). SingleR (42) using the ImmGen compendium (43) identified cell types unique to the immune repertoire and T cells were filtered for this analysis. Seurat R package (44) was used for clustering and differential expression analysis. CD4^+^/CD8^+^ classification was performed using gene expression level of markers (*CD4* or *CD8A*) with a cutoff > 60%. CellChat was used to evaluate the number and relative strengths of interactions between T cells and antigen presenting cells (45).

Geneset variation analysis (GSVA) (46) was applied to SCT-transformed counts against an updated reference gene panel generated from LM22 hematopoietic cells (47). GSVA scores were obtained for each cell and the sub-type with the maximum value for each cell was classified as that sub-type. Differential gene expression (DGE) analysis was carried out via the Model-based Analysis of Single-cell Transcriptomics (MAST) wrapper (48) in Seurat. DGE was applied to clusters specific to either CD4 or CD8 cell-types in a one-vs-all approach and the logFC and p-value ≤0.05 were reported.

Gene Set Enrichment Analysis (GSEA) was performed using the clusterProfiler package to assess enrichment in differentially expressed genes (DEGs) from published and curated immunologic signature gene sets in MSigDB. Parameters were set to include a minimum of 25% expressing cells and a minimum average log2 fold change of 0.25. The FindMarkers function in Seurat, with the MAST wrapper, was used to rank DEGs by decreasing average log2 fold change (-log2FC) and adjusted p-value (-log10). The ranked genes were then input into the GSEA enrichment function using the ImmuneSigDB gene sets from the MSigDB website.

### Statistical Analyses

Statistical tests (Student’s t-test, ANOVA, Non-linear curve fit, Kaplan-Meier test) are indicated in the figure legends. Graphing and statistical analyses were performed in Prism (GraphPad). P or FDR values of <0.05 were considered significant, and error bars represent standard error of the mean (SE). The Broad Institute’s Morpheus software was used to generate heat maps (https://software.broadinstitute.org/morpheus, Broad Institute, Cambridge, MA).

## Results

### Genome-wide analysis of AhR-regulated genes in murine and human LUAD cells

To generate a global transcriptomic profile of AhR-regulated genes in LUAD cells, the AhR was deleted from murine CMT167 (CMT167^AhR-KO^) and human A549 (A549^AhR-KO^) cells by CRISPR/Cas9 gene editing. Controls were transduced with the Cas9-containing vector without guide RNA (CMT167^Cas9^). AhR knockout in two CMT167 clones, D2 and C1, and A549 clones, P15 and P16, was confirmed by western blotting and by failure of a strong AhR ligand, 6-formylindolo(3,2-b)carbazole (FICZ), to induce *CYP1B1*, a canonical AhR-driven gene (**Supplementary Fig. 1A,B,D,E**). Lentivirus transduction had no effect on the *in vitro* growth of AhR-KO lines (Supplementary Fig. 1C,F). Bulk RNA sequencing of CMT167^Cas9^ and CMT167^AhR-KO^ cells demonstrated that 883 genes were significantly downregulated (≥2-fold, FDR<0.05) and 756 genes were significantly upregulated (≥2-fold, FDR<0.05) in both CMT167^AhR-KO^ and A549^AhR-KO^ cells. (The complete list of differentially expressed genes in both lines can be found at GEO accession GSE241980). As expected, expression of the AhR-driven *CYP1A1* and *CYP1B1* genes decreased significantly after AhR knockout in both cell lines (**Fig. 1A,B**).

**Figure 1.**
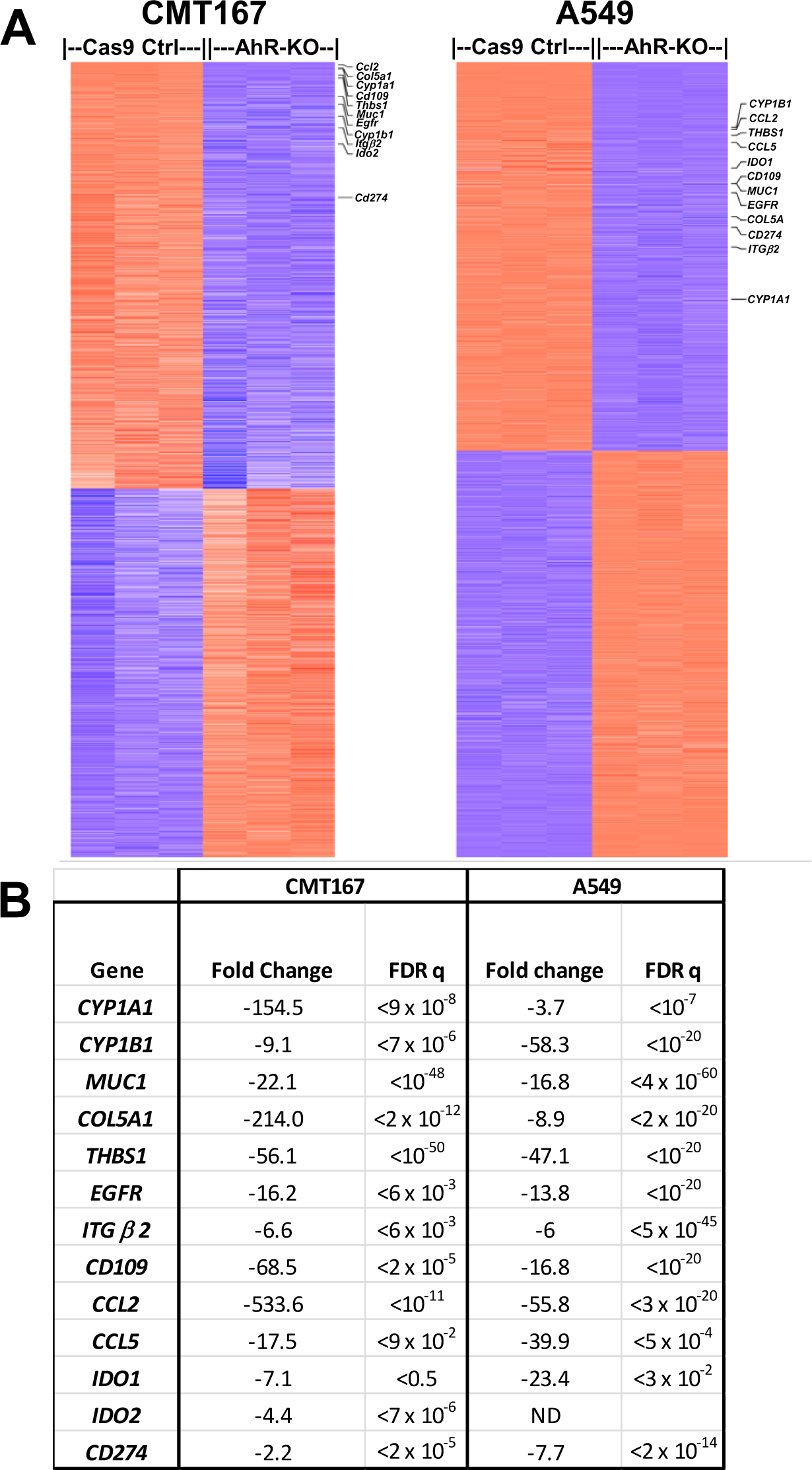
Bulk RNA-seq analysis of AhR-knockout murine and human lung adenocarcinoma cell lines. RNA was extracted from three sets of CMT167^Cas9^ control, CMT167^AhR-KO^, A549^Cas9^ control, or A549^AhR-KO^ cells, reversed transcribed, and cDNA sequenced using the Illumina NextSeq 2000 platform. Data are presented as counts defined as the number of read pairs aligning uniquely to the genome in proper pairs and assigned to a single Ensembl Gene locus for each gene transcript. **A)** Heatmap of all genes with 2-fold or greater change in expression with a false discovery rate (FDR) <0.05 after AhR knockout, as comparison with Cas9 controls, in CMT167 (left) or A549 (right) cells. **B)** Representative cancer- or immune-related genes found to be highly differently downregulated upon AhR knockout in CMT167 and A549 cells.

Included in the set of downregulated genes in both lines were five genes previously shown to be important in LUAD but not commonly associated with the AhR, i.e., *MUC1* (49), *COL5A1* (50), *THBS1* (51), *EGFR* (52), and *ITGβ2* (53) (**Fig. 1A,B**). Of particular note, *ITGβ2* expression on malignant cells is associated with myeloid-derived suppressor cell infiltration in LUAD (54). Six additional genes associated with immune regulation in LUAD, *CD109* (55), *CCL2* (56), *CCL5* (57), *IDO1, IDO2* (58), and *CD274* (*PD-L1*) (59), were significantly downregulated in one or both cell lines. Down-regulation of *Muc1*, *Col5a1*, *Thbs1*, *Egfr*, *Cd109*, *Ccl2* and *Ccl5* in CMT167^AhR-KO^ cells was confirmed by RT-qPCR (**Fig. 2**).

**Figure 2.**
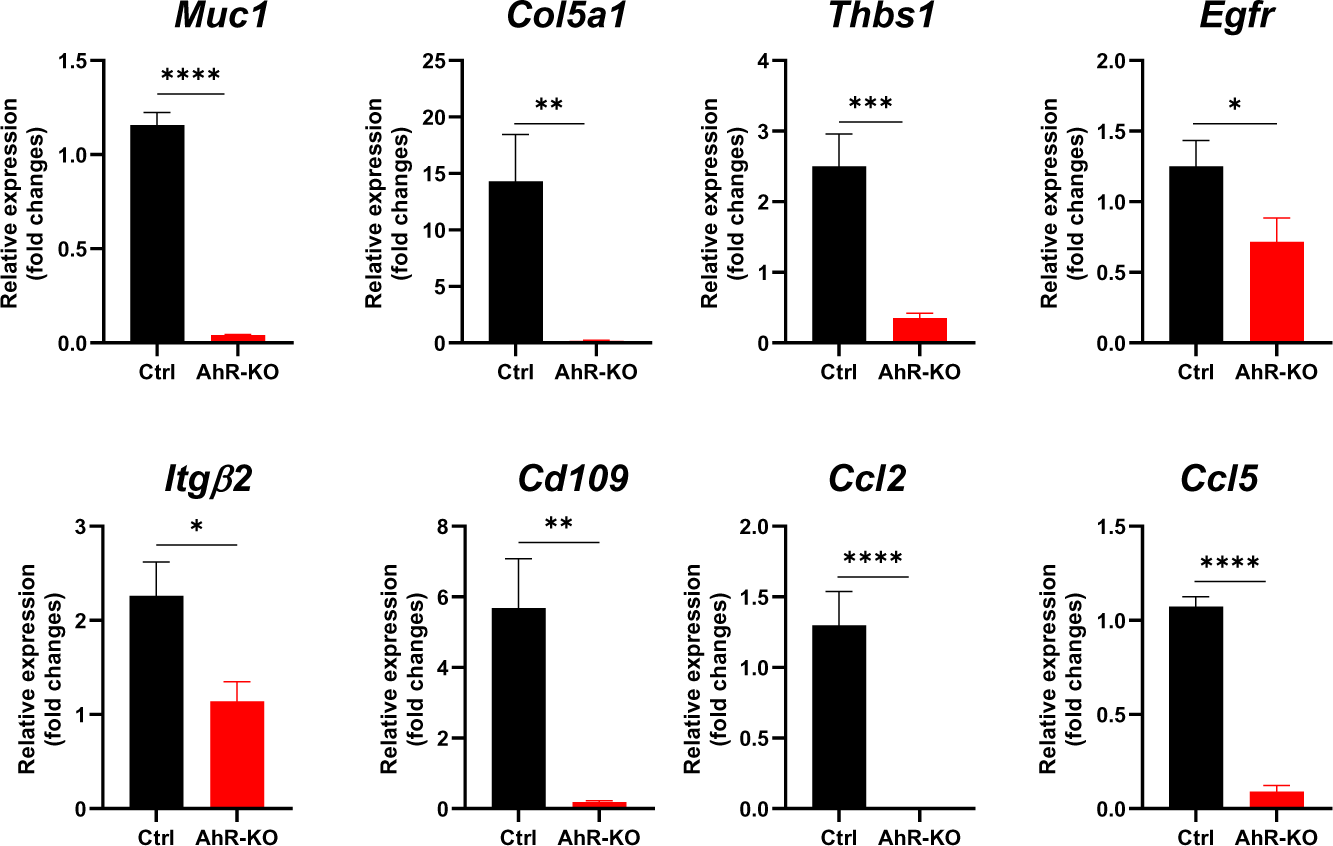
AhR knockout reduces expression of several LUAD-associated genes. Eight genes associated with LUAD and downregulated in CMT167^AhR-KO^, as indicated by RNA-seq (Fig. 1), were quantified by RT-qPCR. There were no statistical differences here or elsewhere between gene levels in AhR^WT^ or AhR^Cas9^ cells. Therefore, results from those two control lines were pooled and referred to here and elsewhere as “Ctrl”. Data are presented as means + SE from three independent experiments with duplicates. in each *p<0.05, **p<0.01, ***p<0.001, ****p<0.0001 (Student’s t-test, equal variance).

The apparent down-regulation of *IDO1 and IDO2* in LUAD cells, as seen by RNA-seq, was consistent with our studies and those of others demonstrating that the AhR regulates IDO1 or IDO2 in breast cancer (27, 60, 61) and dendritic cells (62, 63). Similarly, the down-regulation of *CD274* after AhR knockout is consistent with data showing a correlation between AhR and PD-L1 expression in LUAD patients (35). Given the critical role of IDO1, IDO2 and CD274 in suppression of tumor-specific responses, we then sought to confirm AhR control of these genes in LUAD cells and to assess the relationship of the AhR to other signaling pathways, specifically the IFNγ-induced JAK/STAT pathway, known to influence expression of these immune checkpoints.

### AhR regulates PD-L1 and IDO expression in CMT167 cells

While not yet shown to be effective as monotherapies (64), IDO inhibitors are still in clinical trials for treatment of LUAD, generally in combination with PD-1/PD-L1 blockade (65–67). Given the importance of these mediators of immune suppression and the RNA-seq results suggesting AhR control of *IDO* and *CD274*, we then sought to confirm AhR regulation of *Ido1, Ido2* and *Cd274* in CMT167 cells. RT-qPCR analysis confirmed significant decreases in baseline levels of *Cd274*, *Ido1*, *Ido2*, and, as positive controls, *Cyp1a1*, and *Cyp1b1*, in CMT167^AhR-KO^ cells as compared with control cells (**Fig. 3A, first two bars in each plot**). (*Tdo2* expression was below the level of detection). These results suggest the presence of endogenous AhR ligand(s) that drives baseline levels of these genes. Treatment with 10 µM of the environmentally common smoke constituent and AhR agonist, benzo(a)pyrene (B(a)P), significantly increased expression of all five genes in control cells and those increases were significantly muted or absent in CMT167^AhR-KO^ cells (**Fig. 3A, second two bars in each plot**).

**Figure 3.**
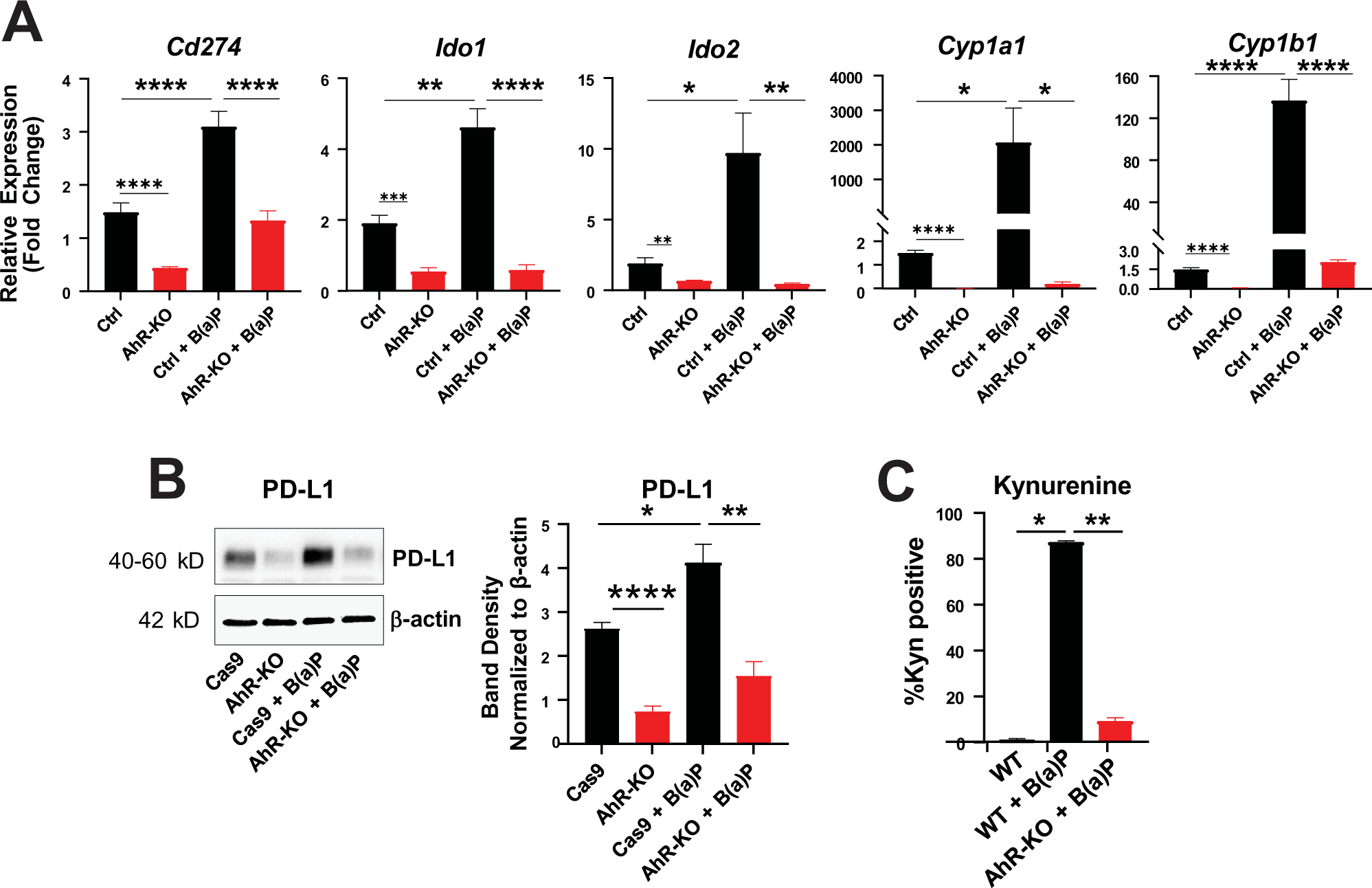
B(a)P, a cigarette smoke constituent, induces PD-L1 and IDO in murine CMT167 cells. **A)** Expression of *Cd274*, *Ido1*, *Ido2*, *Cyp1a1*, and *Cyp1b1* mRNA was quantified by RT-qPCR in control and CMT167^AhR-KO^ cells (left two bars in each plot) or after 72 hours of treatment with 10 µM benzo(a)pyrene (B(a)P)(right two bars in each plot). Data from three independent experiments, each in duplicate or triplicate are presented as *Gapdh*-normalized means + SE. **B)** Protein extracted from cells treated as in (**A**) was probed by western immunoblotting for PD-L1 and, as a loading control, β-actin. One of three representative western blots is shown on the left and β-actin-normalized protein band densities are on the right. Band density data are presented as means from three experiments, each in triplicate, + SE. **C)** The percentage of Kyn^+^ CMT167^WT^ or CMT167^AhR-KO^ cells treated with vehicle or B(a)P as in (**A**) was quantified by flow cytometry. Data from two experiments, each in triplicate, are presented as means + SE from. *p<0.05, **p<0.01, ****p<0.0001 (Student’s t-test, equal variance).

Similarly, naïve CMT167^AhR-KO^ cells expressed significantly less baseline PD-L1 protein than control cells as seen by a representative western immunoblot (**Fig. 3B, left**) and by quantification of β-actin-normalized band densities from three independent experiments (**Fig. 3B, right**). B(a)P increased PD-L1 protein levels and that increase was significantly muted in CMT167^AhR-KO^ cells (**Fig. 3B**). Consistent with AhR control of *Ido1 and Ido2* levels, B(a)P increased baseline levels of Kyn in CMT167^WT^ cells and that increase was significantly lower in CMT167^AhR-KO^ cells (**Fig. 3C**). These results demonstrate that both baseline and environmental chemical-induced levels of two important immune checkpoints PD-L1 and IDO1/2, and a related effector of immunosuppression, Kyn, are AhR controlled.

### IFNγ-mediated induction of PD-L1, IDO1, JAK2, and STAT1 is partially regulated by the AhR

Although most often associated with protective T cell immunity, IFNγ also can upregulate PD-L1 (12, 13) and IDO (68) thereby suggesting the counterintuitive conclusion that chronic production of IFNγ, presumably by TILs, may be counterproductive to the anti-tumor immune response. To determine if the AhR plays a role in IFNγ-mediated induction of these immune checkpoints in LUAD cells, control and CMT167^AhR-KO^ cells were treated with IFNγ and *Cd274*, *Ido1*, and *Ido2* mRNA quantified by RT-qPCR after 24 (*Cd274*) or 72 (*Ido1*, *Ido2*) hours. *Cyp1a1* and *Cyp1b1* mRNA levels were measured as controls. As shown previously (**Fig. 3**), CMT167^AhR-KO^ cells expressed significantly lower baseline levels of all five genes than control cells (**Fig. 4A**, **first two bars in each plot**). In control cells, IFNγ induced about a 95-, 15,500-, and 60-fold increase in *Cd274*, *Ido1*, and *Ido2* expression, respectively (**Fig. 4A, third bar**). However, IFNγ-mediated induction of all three genes was significantly muted in CMT167^AhR-KO^ cells (**Fig. 4A, fourth bar**). As expected from the IFNγ-driven increase in *Ido1 and Ido2*, generators of tryptophan-derived AhR ligands, both *Cyp1a1* and *Cyp1b1* transcripts increased in an AhR-dependent way (**Fig. 4A**). To our knowledge, this is the first demonstration, in any context, of this critical T cell cytokine increasing these two AhR-driven metabolic gene transcripts.

**Figure 4.**
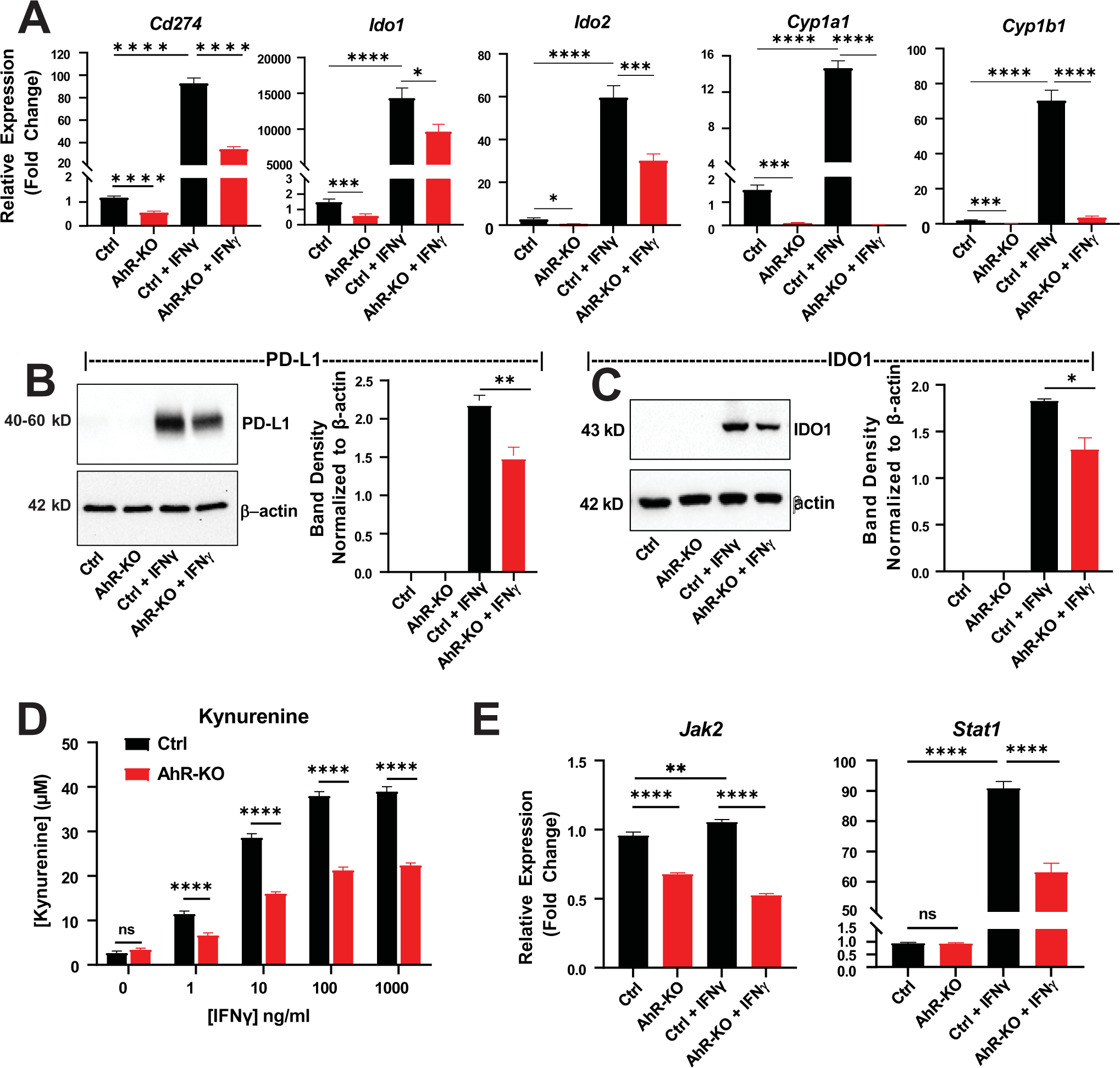
The AhR mediates IFNγ induction of immune-related genes *Cd274, Ido1/2, Jak2,* and *Stat1* in murine LUAD CMT167 cells. **A)** CMT167^Ctrl^ or CMT167^AhR-KO^ cells were untreated or treated with 100 ng/ml IFNγ. *Cd274* mRNA was quantified by RT-qPCR 24h later. *Ido1*, *Ido2*, *Cyp1a1*, and *Cyp1b1* mRNA was quantified 72h later. Data are from three independent experiments, each in duplicate or triplicate, are expressed as fold change of *Gapdh*-normalized means + SE. **B)** CMT167^Ctrl^ or CMT167^AhR-KO^ cells were left untreated or treated for 24h with 100 ng/ml IFNγ and PD-L1 protein expression assayed by western immunoblotting. A representative immunoblot is on the left and β-actin normalized band densities, averaged from three independent experiments, is on the right. (Bands from IFNγ-treated cells reached saturation prior to bands from untreated cells becoming visible). **C)** CMT167^Ctrl^ or CMT167^AhR-KO^ cells were treated for 24h with IFNγ and IDO1 protein expression assayed by western immunoblotting. A representative immunoblot is on the left and β-actin normalized band densities, averaged from three independent experiments, are on the right. **D)** CMT167^Ctrl^ or CMT167^AhR-KO^ cells were treated with 1-1000 ng/ml IFNγ for 24h and Kyn released into the media quantified via colorimetric assay. Data are averaged from two independent experiments each in quadruplicate + SE. **E)** CMT167^Ctrl^ or CMT167^AhR-KO^ cells were treated with 100 ng/ml IFNγ and *Jak2* and *Stat1* mRNA quantified 24h later. RT-qPCR data are from three independent experiments, each in triplicate, and presented as *Gapdh*-normalized means + SE. *p<0.05, **p<0.01, ***p<0.001, ****p<0.0001 (Student’s t-test, equal variance).

As expected from RT-qPCR results, IFNγ-mediated induction of PD-L1 (**Fig. 4B**) and IDO1 (**Fig. 4C**) protein was significantly lower in CMT167^AhR-KO^ cells. (Note that IFNγ led to such dramatic increases in PD-L1 and IDO1 protein that the exposure time in the Western blots had to be shortened and, therefore, was insufficient for revealing baseline PD-L1 or IDO1 levels). Consistent with AhR-regulation of IFNγ-induction of IDO, increasing IFNγ doses increased Kyn levels, as quantified by a Kyn-dependent colorimetric assay, in a dose-dependent manner and that increase was reduced in CMT167^AhR-KO^ cells (p<0.0001) (**Fig. 4D**).

Previous studies demonstrated a role for the JAK/STAT signaling pathway in PD-L1 expression in lung cancer (69) and a correlation between JAK/STAT, IDO, and PD-L1 in other cancers (70–72). The JAK/STAT pathway can be activated by IFNγ in various cancers including LUAD (73). Therefore, a possible role for the AhR in regulating IFNγ-mediated upregulation of *Jak* or *Stat* was evaluated. Baseline *Jak2* levels, as quantified by RT-qPCR, were lower in CMT167^AhR-KO^ cells relative to controls (**Fig. 4E, left, first two bars**). IFNγ modestly but significantly increased *Jak2* in control cells but not in CMT167^AhR-KO^ cells. No differences in baseline *Stat1* levels were seen between control and CMT167^AhR-KO^ cells. However, IFNγ increased *Stat1* levels by ∼90-fold in control cells and this *Stat1* induction was significantly reduced in CMT167^AhR-KO^ cells (**Fig. 4E, right**) implicating the AhR in IFNγ induction of *Jak2* and *Stat1*.

These studies were then extended to the human A549 LUAD line with similar, if not more profound results. In A549 cells, AhR knockout significantly reduced baseline levels of *CD274*, *IDO1,* and *CYP1B1* mRNAs (**Fig. 5A, first two bars in each graph**). (*IDO2* and *CYP1A1* mRNAs in A549 cells were at or below the level of detectability). IFNγ significantly increased expression of all three genes in control cells but that increase was significantly muted or absent in A549^AhR-KO^ cells (**Fig. 5A, second two bars**). As expected from the *CD274* RT-qPCR results, the baseline percentages of PD-L1^+^ cells, as quantified by flow cytometry, were significantly reduced in A549^AhR-KO^ cells (**Fig. 5B, first two bars**). IFNγ increased the percentage of PD-L1^+^ A549^Ctrl^ cells but not the percentage of PD-L1^+^ A549^AhR-KO^ cells (**Fig. 5B, second two bars**).

**Figure 5.**
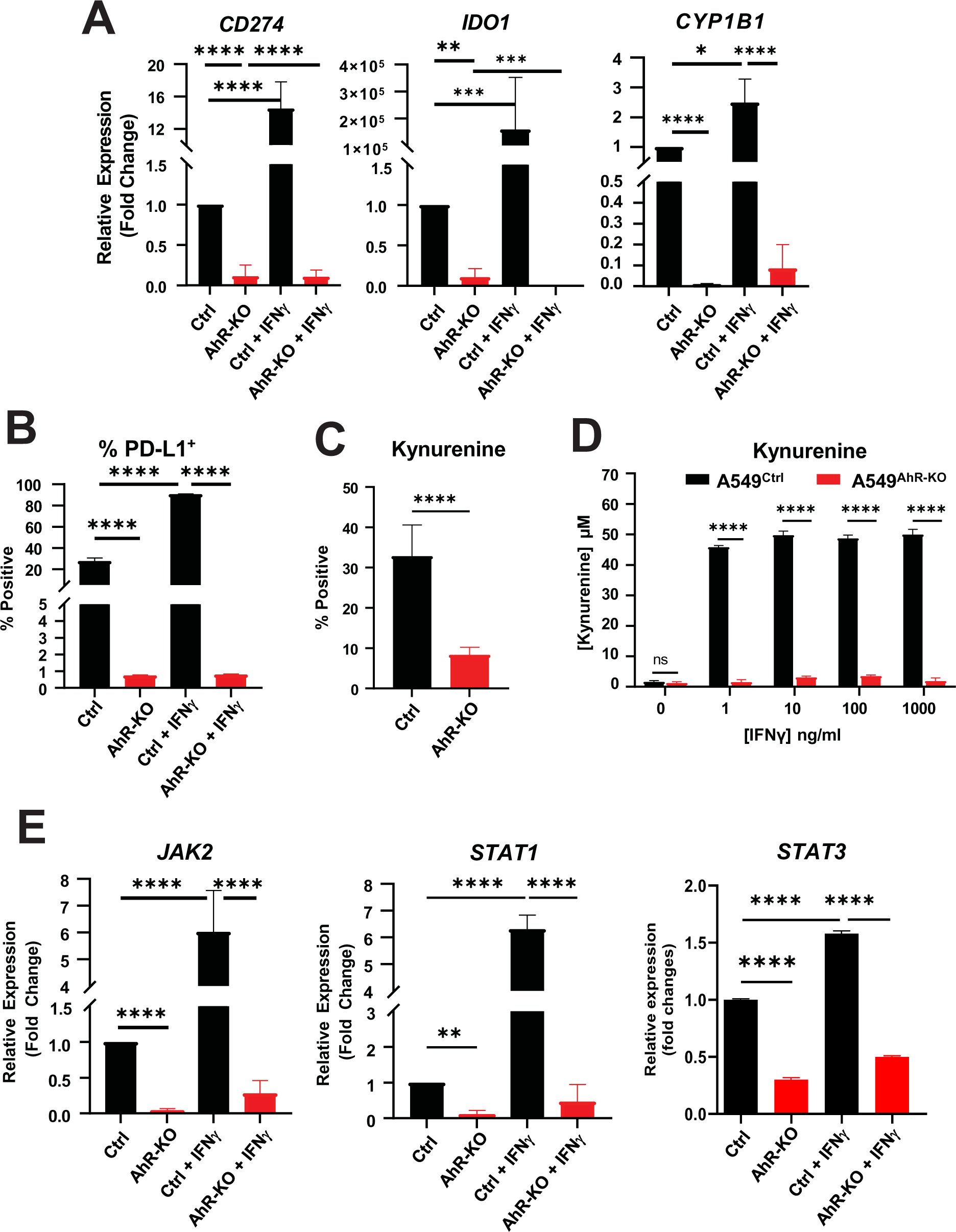
The AhR mediates IFNγ induction of immune-related genes *CD274, IDO1, JAK2, STAT1*, and *STAT3* in human LUAD A549 cells. **A)** A549^Ctrl^ or A549^AhR-KO^ cells were untreated or treated with 100 ng/ml IFNγ for 24h and *CD274*, *IDO1,* and *CYP1B1* expression quantified by RT-qPCR. Data from four experiments, each in triplicate, are represented as fold change of *GAPDH*-normalized means + SE. **B)** The percent positive PD-L1^+^ cells treated as in (**A**) was quantified by flow cytometry. Data from three experiments, each in triplicate, are presented as mean percent PD-L1^+^ + SE. **C)** The baseline percent of Kyn^+^ A549^ctrl^ and A549^AhR-KO^ cells in two experiments, each in triplicate, was determined by flow cytometry. **D)** A549^Ctrl^ or A549^AhR-KO^ cells were treated with 0-1000 ng/ml IFNγ for 24h and Kyn release quantified by the Kyn-specific colorimetric assay using a standard Kyn curve. Data from two experiments, each in quadruplicate, are presented as average μM Kyn + SE. **E)** A549^Ctrl^ or A549^AhR-KO^ cells were left untreated or treated with 100 ng/ml IFNγ for 24h and baseline or 100 ng/ml IFNγ-induced *JAK2*, *STAT1,* and *STAT3* expression quantified by RT-qPCR. Data from four experiments, each in triplicate, are presented as average fold change of *Gapdh*-normalized means + SE. *p<0.05, **p<0.01, ***p<0.001, ****p<0.0001 (Student’s t-test, equal variance).

As expected from the IFNγ-induced *IDO1* mRNA quantification (**Fig. 5A**), AhR knockout significantly decreased the percentage of Kyn^+^ cells (**Fig. 5C**). Similarly, titered concentrations (1-1000 ng/ml) of IFNγ significantly induced Kyn production in control cells, but minimally if at all in A549^AhR-KO^ cells (**Fig. 5D**).

Consistent with data obtained in murine CMT167 lines (**Fig. 4**), baseline *JAK2*, *STAT1,* and *STAT3* mRNA levels were significantly lower in A549^AhR-KO^ cells as compared with controls (**Fig. 5E, first two bars**). IFNγ significantly increased *JAK2*, *STAT1*, and *STAT3* in control but not in A549^AhR-KO^ cells (**Fig. 5E, third and fourth bars**). IFNγ did not affect AhR levels in either CMT167 or A549 cells (not shown).

Collectively, the data generated in murine CMT167 and human A549 LUAD cells demonstrate that the AhR influences the baseline and IFNγ-induced expression of key immune regulators, IDO and PD-L1, as well as components of the JAK/STAT signaling pathway that are known to regulate PD-L1 and IDO expression.

### AhR deletion in CMT167 cells imparts partial immune protection *in vivo*

The lower levels of baseline and IFNγ-induced IDO1/2 and PD-L1 in CMT167^AhR-KO^ cells suggest the potential for an enhanced immune response to these cells *in vivo*. To test this hypothesis, growth of wildtype, Cas9 control and AhR-KO CMT167 clones C1 and D2 cells was determined in syngeneic C57BL/6 mice. CMT167^WT^ and CMT^Cas9^ tumors emerged at approximately day 14 in each of the 16 mice injected and grew rapidly over the next 10 days (**Fig. 6A**). All of these mice required euthanasia by day 30 because of skin lesions over the tumor cell injection site. In contrast, of the 64 mice injected with CMT167^AhR-KO^ clones C1 and D2, 49 (77%) failed to grow tumors by day 53 (**Fig. 6A, red arrow**). Of the CMT167^AhR-KO^ tumors that did grow, they grew at a significantly slower pace than control tumors (**Fig. 6B**).

**Figure 6.**
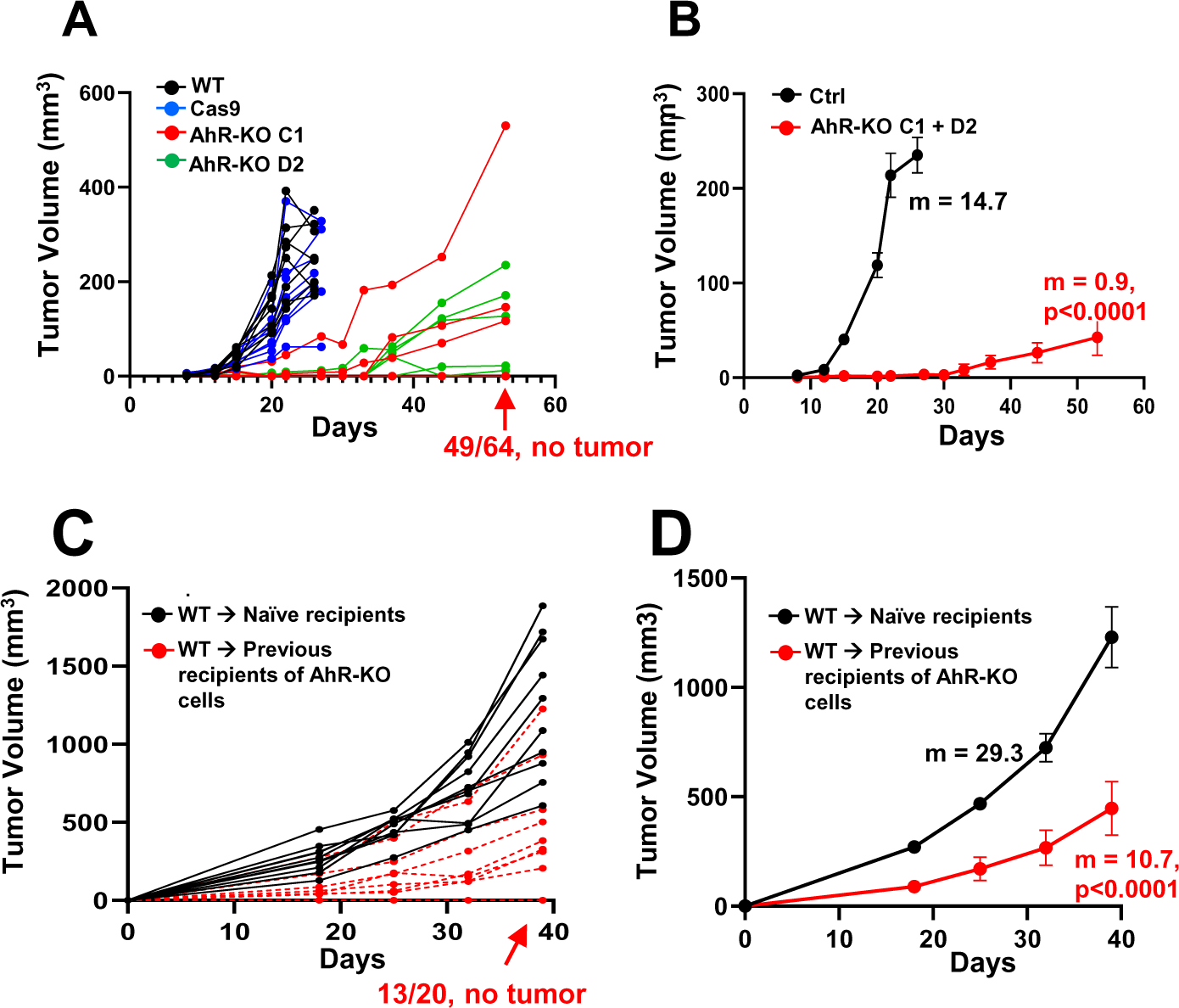
AhR deletion in CMT167 cells leads to decreased tumor burden and resistance to re-challenge with wildtype cells. **A)** 10^6^ CMT167^WT^ (black lines), CMT167^Cas9^ (blue lines), CMT167^AhR-KO^ clone C1 (red lines), or CMT167^AhR-KO^ clone D2 (green lines) cells were injected subcutaneously into syngeneic C57BL/6 mice and tumor growth determined over a 53-day period. 100% of mice inoculated with CMT167^WT^ cells grew tumors. No tumors were detected in 49 nine of 64 possible CMT167^AhR-KO^ tumors (77%)(red arrow). **B)** Growth curves of control (CMT167^WT^ + CMT^Cas9^, n=32) and CMT167^AhR-KO^ (clones C1+D2, n=15) tumors from “**A**” were averaged. Data are presented as means ± SE, m = slopes. p<0.0001 by non-linear best fit curve comparison. **C)** Mice that had been inoculated 65 days earlier with CMT167^AhR-KO^ cells (n=10 CMT167^AhR-KO^ clone C1, n=10 CMT167^AhR-KO^ clone D2) and which had not grown tumors were re-challenged with CMT167^WT^ cells and tumor growth tracked over 40 days (red dashed lines). Naïve age-matched mice (n=20) were injected with CMT167^WT^ cells as positive controls (black lines). 100% of naïve mice inoculated with CMT167^WT^ cells grew tumors. No tumors were detected in 13 of 20 mice (65%) injected 65 days previously with CMT167^AhR-KO^ cells (red arrow). **D)** Growth curves of CMT167^WT^ tumors injected in naïve mice or into previous recipients of CMT167^AhR-KO^ cells were averaged. Data are presented as means + SE. m=slope by linear regression, p<0.0001.

To determine if the slow/lack of growth of CMT167^AhR-KO^ tumors reflected a heightened immune response to CMT167^AhR-KO^ cells as compared with controls, 20 of the mice that failed to generate CMT167^AhR-KO^ tumors by day 53 were inoculated in the contralateral flank with CMT167^WT^ cells. Ten age-matched naïve mice were injected with CMT167^WT^ cells as positive controls. Of the 20 mice that had previously been inoculated with CMT167^AhR-KO^ cells, none grew tumors at the original site of CMT167^AhR-KO^ cell inoculation within 40 days of the rechallenge and 13 of the 20 (65%) never grew wildtype tumors in the contralateral flank (**Fig. 6C, red arrow**). The seven CMT167^WT^ tumors that did grow grew significantly more slowly than CMT167^WT^ tumors generated in naïve controls (**Fig. 6D)**. These data indicate that AhR deletion in CMT167 cells induces a systemic and relatively long-lasting immunity with the potential for complete tumor clearance.

### AhR deletion in CMT167 cells enables tumor-infiltrating T cell recruitment

To characterize the nature of the immunity imparted by transplantation of AhR-knockout CMT167 cells, CMT167^WT^ and CMT167^AhR-KO^ tumors were excised, formalin fixed and sectioned five weeks after transplantation and evaluated by immunofluorescence for AhR expression and infiltration of CD45^+^ cells. While AhR^+^ CMT167^WT^ tumors were nearly devoid of immune cells, significant numbers of CD45^+^ cells were seen in CMT167^AhR-KO^ tumors (**Fig. 7A**). When quantified, this translated to a ∼5-fold higher density of CD45^+^ cells/mm^3^ in the CMT167^AhR-KO^ tumors as compared with CMT167^WT^ cells (p<0.0001)(**Fig. 7B)**.

**Figure 7.**
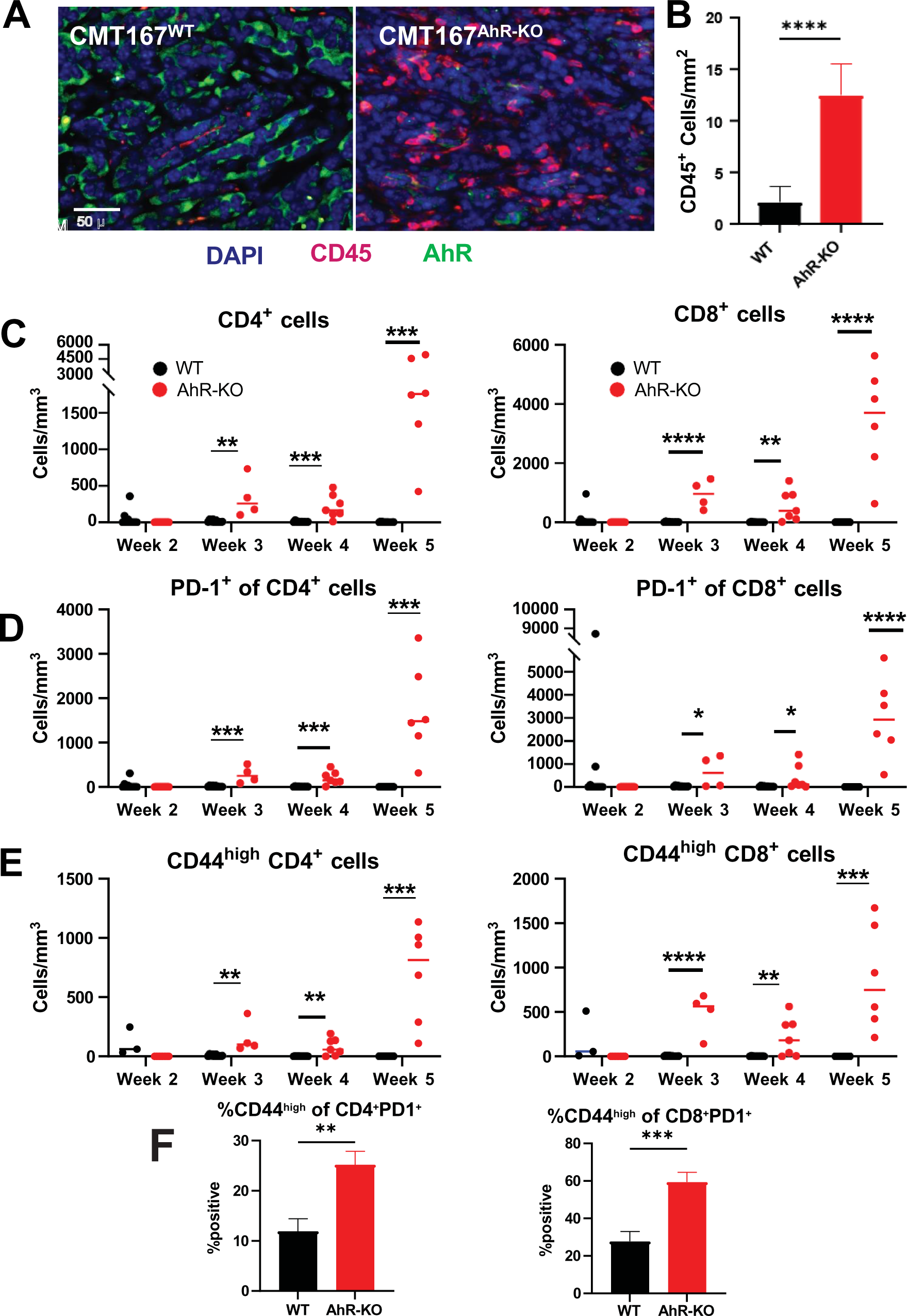
CMT167^AhR-KO^ tumors have a higher density of infiltrating CD4^+^ and CD8^+^ T cells than CMT167^WT^ tumors. 10^6^ CMT167^WT^ or CMT167^AhR-KO^ cells were injected subcutaneously into syngeneic C57BL/6 mice. Tumors, if present, were excised between two and five weeks after cell injection. Approximately half of each five-week tumor was fixed and sectioned for immunofluorescent studies and the remaining tumor half digested to recover infiltrating leukocytes. **A)** Representative immunofluorescent images from a total of five CMT167^WT^ and four CMT167^AhR-KO^ five-week tumors (three sections/tumor) stained with DAPI (blue), AhR-specific antibody (green), and CD45-specific antibody (red) are shown. **B)** Mean density (number of cells/tumor mm^2^) + SE of CD45^+^ cells from five CMT167^WT^ and four CMT167^AhR-KO^ five-week tumors. **C-E**) Tumor infiltrating cells were recovered, counted, and stained for CD45, CD4, CD8, PD-1, and CD44 and analyzed by flow cytometry. Each dot represents the number of cells/mm^3^ (i.e. cell density) from one tumor. Data were obtained from two to 10 mice per group per week depending on tumors available to excise. *p<0.05, **p<0.01, ***p<0.001, ****p<0.0001 (Multiple comparisons t tests). **F)**. The percent of all CD4^+^PD-1^+^ (left) or CD8^+^PD-1^+^ (right) T cells that are also CD44^high^. Data from two independent experiments, six mice/condition/experiment, are presented as the percent positive + SE. **p<0.01, p<0.001 (Student’s t-test, equal variance).

Flow cytometric analysis of CD45^+^ immune cells from tumors that were excised and digested at two, three, four, and five weeks after transplantation revealed relatively few CD45^+^CD4^+^ or CD45^+^CD8^+^ T cells in CMT167^WT^ tumors, resulting in a very low T cell density at any time point (**Fig. 7C, black circles**). (Representative dot plots are provided in **Supplemental Fig. 2**). In contrast, an increasing density of CD4^+^ and CD8^+^ T cells was noted over time in CMT167^AhR-KO^ tumors (**Fig. 7C**, **red circles**). Indeed, the smaller CMT167^AhR-KO^ tumors generally had a greater absolute number of infiltrating CD4^+^ and CD8^+^ T cells than the larger CMT167^WT^ tumors (**Supplemental Fig. 3A).** While IFNγ-producing T cells were seen in CMT^WT^ tumors, a significantly greater density of IFNγ-producing CD4^+^ and CD8^+^ T cells was seen in CMT167^AhR-KO^ tumors (**Supplemental Fig. 3B).**

Corresponding to this increase in total CD4^+^ and CD8^+^ T cells, the density of CD4^+^PD-1^+^ or CD8^+^PD-1^+^ T cells increased over time in the CMT167^AhR-KO^ but not in control tumors (**Fig. 7D**). Although associated with T cell exhaustion, high PD-1 levels on T cells have also been positively associated with LUAD survival after anti-PD-1 therapy (74). In addition, PD-1 has recently been shown to mark a subset of tumor neoantigen-specific tissue-resident memory T cells in LUAD (74), HNSCC (75), colorectal cancer (75) and sarcoma (76). In this vein, the density of CD4^+^ or CD8^+^ T cells expressing a CD44^high^ activated/memory phenotype increased over time in the CMT167^AhR-KO^ but not in control tumors (**Fig. 7E**). Furthermore, the percentages of CD4^+^PD-1^+^ or CD8^+^PD-1^+^ cells that were also CD44^high^ were significantly higher in CMT167^AhR-KO^ tumors (**Fig. 7F**). These data suggest that tumor-infiltrating T cells in the CMT167^AhR-KO^ tumors express more of an activation/memory phenotype than those from wildtype tumors.

Finally, we note that 77% of the CMT167^AhR-KO^ cell-transplanted mice never grew tumors and therefore were not evaluable for T cell infiltration. Therefore, analysis of only the 23% of CMT167^AhR-KO^ tumors that did form may underestimate the extent of T cell recruitment during complete rejection of AhR-KO tumors.

### Single cell RNA-sequencing analysis of tumor-infiltrating T cells

To more definitively categorize tumor infiltrating T cell subsets, scRNA-seq was performed on sorted CD45^+^ immune cells (>90% viable) from digested 5-week tumors. SingleR (42) and the ImmGen compendium (43) were used to identify T cell subtypes and the Seurat R package (44) was used for clustering and differential gene expression (DGE) analysis. Clustering of CD3 T cells resulted in 16 unique T cell clusters (#0-15) visualized in UMAP plots (**Fig. 8A**). Five clusters were categorized as CD4 T cells (**Fig. 8B green dots; Fig. 8C left**) and 11 were categorized as CD8 T cells (**Fig. 8B purple dots; Fig. 8D right**).

**Figure 8.**
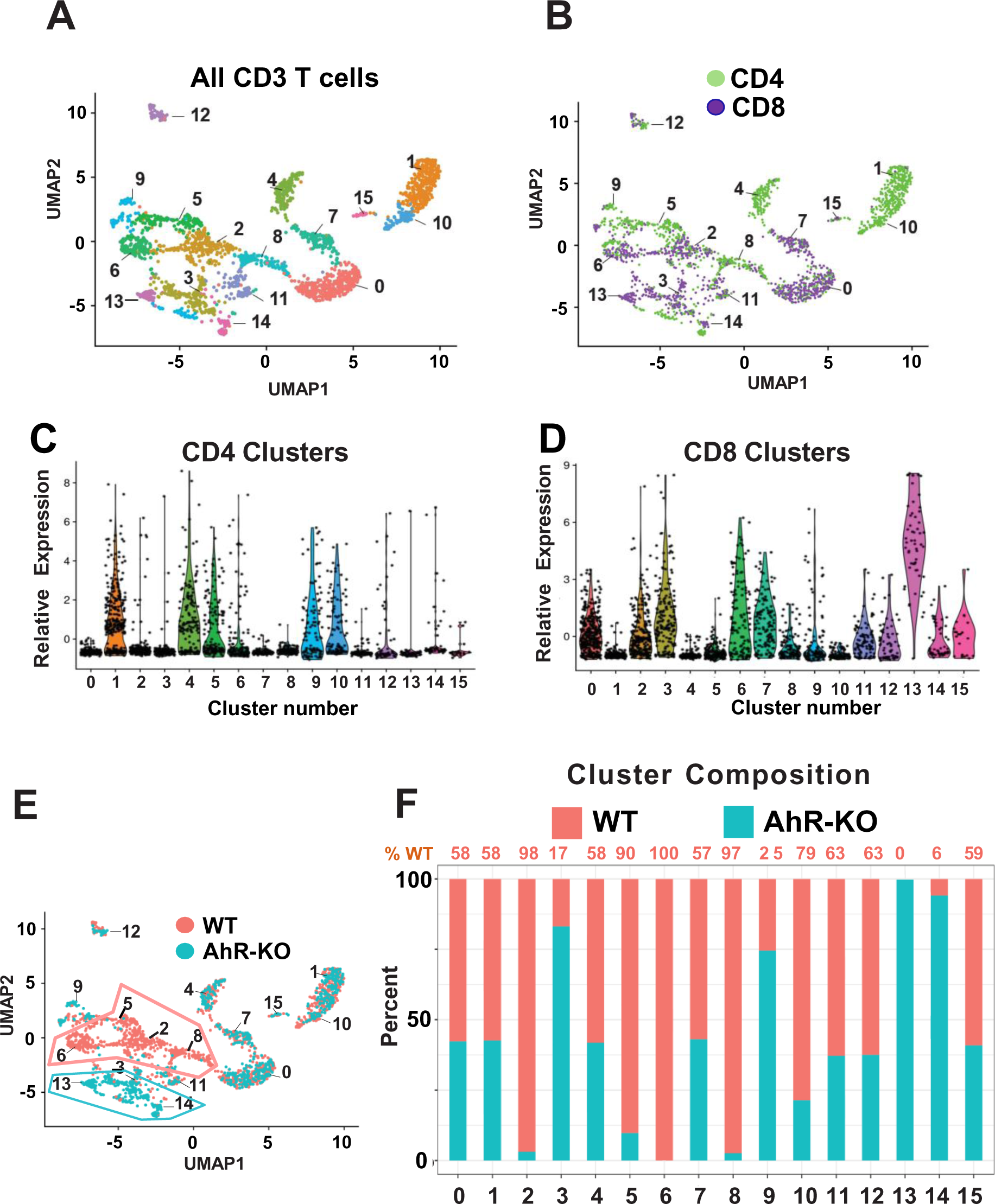
scRNA-seq of CD45^+^ TILs from CMT167^WT^ or CMT167^AhR-KO^ five-week tumors reveals differences in TIL composition. **A)** CMT167^WT^ or CMT167^AhR-KO^ tumors excised five weeks after transplantation as in Fig. 7 were digested and sorted by flow cytometry for CD45^+^ cells. RNA from single cells was then sequenced. Greater than 2000 *CD3^high^* T cells were recovered from each sample. Sixteen unique CD3 Seurat clusters (#0-15) were identified using the Immunological Genome Project (ImmGen) reference compendium (43) and the singleR annotation method. **B)** Clusters were overlayed in green and purple to designate CD4 and CD8 cells respectively. **C,D**) Violin plots identify distinct CD4 (**C**) and CD8 (**D**) T cell populations. **E)** Clusters were overlayed burnt orange or teal to designate cells from CMT167^WT^ or CMT167^AhR-KO^ tumors, respectively. The orange polygon indicates the relative transcriptomic resemblance of clusters 2, 5, 6, 8 from CMT167^WT^ tumors and the teal polygon indicates the relative transcriptomic similarity of clusters 13 and 14 from CMT^AhR-KO^ tumors. **F)** Proportion of cells originating from CMT167^WT^ (orange) and CMT167^AhR-KO^ (teal) tumors within each Seurat cluster. The exact percentage of T cells from CMT167^WT^ tumors is presented at the top.

Quantifying the percentage of CD4 and CD8 T cells from CMT167^WT^ or CMT167^AhR-KO^ tumors that are represented in each of the 16 CD3 clusters revealed that cluster 5 (CD4) and clusters 2, 6 and 8 (CD8) consisted predominantly (≥90%) of T cell subsets from CMT167^WT^ tumors (**Fig. 8E burnt orange polygon, Fig. 8F**) whereas CD8 clusters 13 and 14 consisted predominantly (≥94%) of T cells from CMT167^AhR-KO^ tumors (**Fig. 8E teal polygon, Fig 8F**). Notably, cluster 6 was unique to CD8 T cells from CMT167^WT^ tumors while cluster 13, which expressed the highest level of *CD8a* (**Fig. 8D)**, was unique to T cells from CMT167^AhR-KO^ tumors (**Fig. 8F**).

Gene set variation analysis (GSVA), a rank-based gene-set enrichment method used to assess relative T cell activity [111], indicated that CD4 and CD8 T cells from CMT167^AhR-KO^ tumors were significantly (p<10^-15^ and p<10^-11^ respectively) more active than those from CMT167^WT^ tumors (**Fig 9A**). Pooling differentially expressed genes in CD8 clusters that were comprised of >94% T cells from either CMT167^WT^ or CMT167^AhR-KO^ tumors demonstrated that the aggregate of CD8 clusters 13+14 from CMT167^AhR-KO^ tumors was significantly more active than the aggregate of CD8 clusters 2+6+8 from CMT167^WT^ tumors (**Fig. 9B, left**). The difference in relative activity was even greater when comparing only the clusters that were 100% unique to AhR-KO and wildtype tumors, i.e., cluster 6 (CMT167^WT^) *vs* cluster 13 (CMT167^AhR-KO^)(**Fig. 9B, right**).

**Figure 9.**
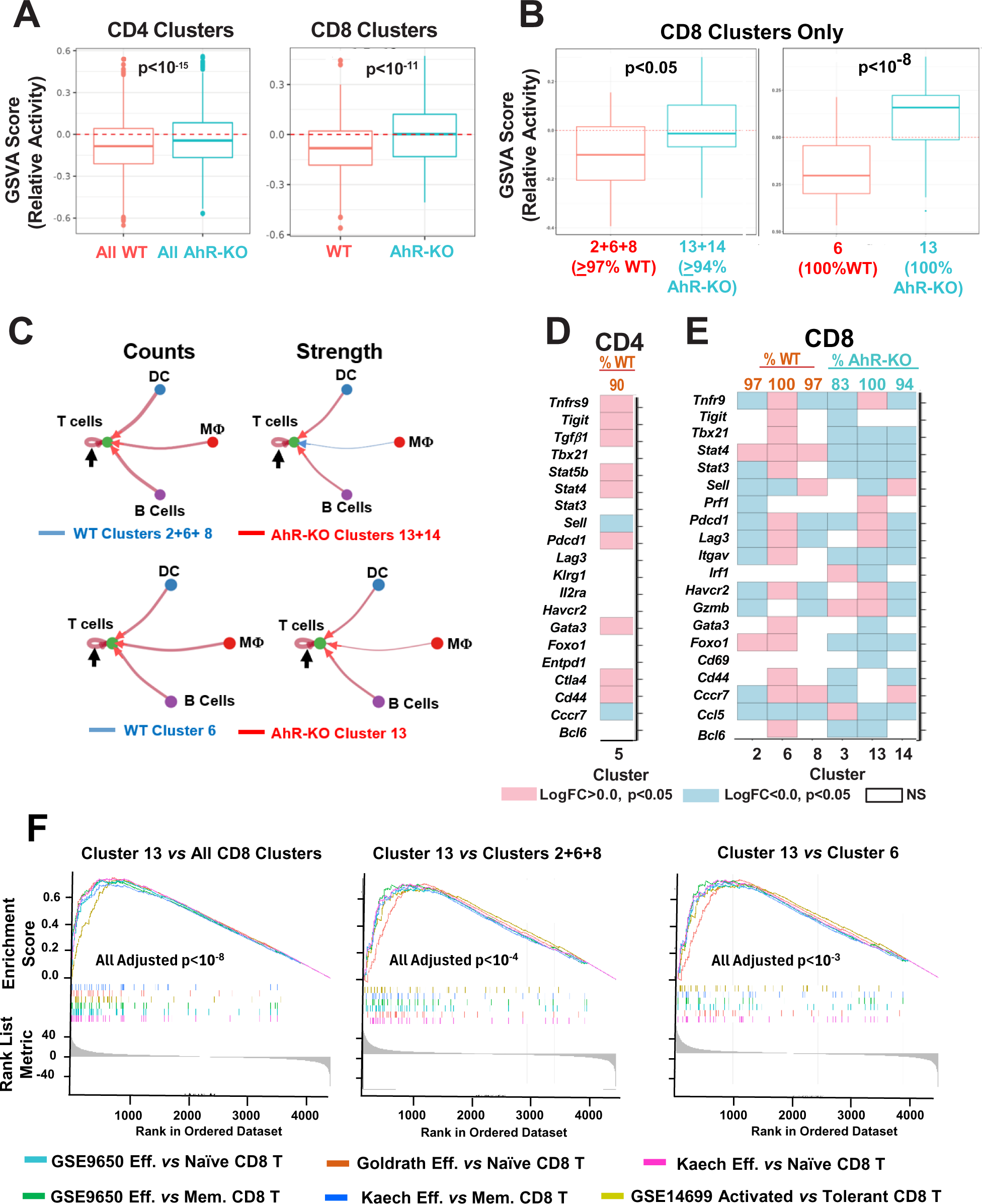
Analysis of T cell clusters infiltrating CMT167^WT^ and CMT167^AhR-KO^ tumors. **A)** GSVA enrichment scores of functional capabilities in all CD4 (left) or CD8 (right) T cell clusters from CMT167^WT^ (orange) and CMT167^AhR-KO^ (teal) tumors. **B) Left:** GSVA enrichment scores of functional capabilities of the aggregate of CMT167^WT^ clusters 2+6+8 vs CMT167^AhR-KO^ clusters 13+14. **Right:** GSVA enrichment scores of functional capabilities in CMT167^WT^ cluster 6 vs CMT167^AhR-KO^ cluster 13. **C)** The CellChat cell-cell communication platform was used to predict the number (left) and strength (right) of incoming signals from dendritic cells (DC), macrophages (MΦ) and B cells to specific CD8 T cell clusters, each of which comprises >90% of the T cells from the respective CMT167^WT^ or CMT167^AhR-KO^ tumors. The color of the lines represents the tumor source of T cells (red = CMT167^AhR-KO^; blue = CMT167^WT^ T cells) and the thickness represents the relative number or strength of incoming signals from APC. **Top)** Incoming signals from APC to the aggregate of CMT167^WT^ clusters 2, 6, and 8 versus incoming signals from APC to the aggregate of CMT167^AhR-KO^ clusters 13 and 14. **Bottom)** Incoming signals from APC to the CMT167^WT^ clusters 6 versus incoming signals from APC to CMT^AhR-KO^ cluster 13. **D, E**) MAST (48) was used to analyze immune-related DEGs (p<0.05) between T cell clusters in which >80% of the cells were derived from CMT167^WT^ or CMT ^AhR-KO^ tumors. **D**) DEGs in CMT167^WT^ CD4 cluster 5 as compared with all other clusters. **E**) DEGs in CMT167^WT^ CD8 clusters 2,6, and 8 and CMT167^AhR-KO^ clusters 3, 13, and 14 as compared with all other clusters. The percentage of cells originating from CMT167^WT^ (orange) or CMT^AhR-KO^ (teal) tumors is restated at the top of the heat maps. **F**) Upregulated genes in CMT167^AhR-KO^ CD8 cluster 13, relative to genes in: 1) all CD8 clusters (left), 2) the aggregate of CMT167^WT^ CD8 clusters 2+6+8 (middle), and 3) CMT167^WT^ CD8 cluster 6 (right) were determined. GSEA was then used to determine enrichment of these three upregulated cluster 13 gene sets in six sets of published upregulated genes from: 1) activated CD8 T cells as compared with naïve CD8 T cells (three gene sets: GSE9650 Eff. vs Naïve CD8 T cells, Goldrath Eff. vs Naïve CD8 T cells, Kaech Eff. vs Naïve CD8 T cells), 2) activated CD8 T cells as compared with memory CD8 T cells (two gene sets: GSE9650 Eff. vs Mem. CD8 T cells, Kaech Eff. Vs Mem. CD8 T cells), and 3) activated CD8 T cells as compared with tolerant CD8 T cells (one gene set: GSE14699 Activated vs Tolerant CD8 T cells).

One type of cell-cell communication that could account for the greater activity seen in CD8 T cell clusters from CMT167^AhR-KO^ tumors would involve interactions of T cells with antigen presenting cells (APC). Therefore, the CellChat cell-cell communication platform (45) was used to estimate the number and strength (weight of cell interactions based on ligand-receptor binding strength/probability) of incoming signaling from dendritic cells (DC), macrophages (MΦ), and B cells to CD8 clusters, with a focus on clusters that represent >90% of T cells from CMT167^WT^ or CMT167^AhR-KO^ tumors. CellChat estimated significantly more incoming interactions (counts) from all three types of APC to the CMT167^AhR-KO^ CD8 clusters 13+14 than from APC to the CMT167^WT^ CD8 clusters 2+6+8 (**Fig. 9C, top left, red lines**). (The color of the lines represents the tumor source of T cells and the thickness represents the relative number of incoming signals from APC). The number of T-T cell interactions in clusters 13+14 was also greater than T cell interactions in clusters 2+6+8 (**Fig. 9C, black arrows**). Similarly, the strength of incoming signals from DC or B cells and between T cells was greater for clusters 13+14 than for clusters 2+6+8 (**Fig. 9C, top right, red lines**). Relatively weak interactions between MΦ and CMT167^WT^ clusters 2+6+8 were noted (**Fig. 9C, top right, blue line**). When comparing cluster 6 and 13, 100% CMT167^AhR-KO^-derived cluster 13 had significantly more and stronger interactions with all three APC than 100% CMT167^WT^-derived cluster 6 (**Fig. 9C, bottom)**.

To assess patterns of single gene expression that could point to cell function, particularly for cluster 13, one (cluster)-vs-all differential analyses were performed considering only CD4 or CD8 clusters with ≥90% representation from either wildtype or AhR-KO tumors. In particular we assessed the relative expression of genes associated with T cell exhaustion/activation and CD8^+^ T cell killing activity. CD4 cluster 5, 90% of which consisted of T cells from CMT167^WT^ tumors, expressed relatively high levels of *Tnfrs9* (CD137)(**Fig. 9D**), a marker which predicts poorer survival in LUAD (77). Cluster 5 also expressed relatively high levels of *Tigit*, *Tgfβ*, *Pdcd1 (Pd-1))*, and *Ctla4*, a phenotype consistent with immunosuppressive or exhausted T cells (78). Among the CD8 clusters (**Fig. 9E**), cluster 6, which is entirely composed of cells from wildtype tumors, was the only CD8 cluster that expressed elevated levels of *Itgav*, *Tnfsrf9*, *Tigit*, *Pdcd-1*, *Lag3*, and *Tim3* (*Havcr2*), all of which are associated with exhausted or immunosuppressive CD8^+^ T cells (78–80). Conversely, the two clusters expressing the highest levels of granzyme B (*Gzmb*), clusters 3 and 13, were composed of T cells predominantly (cluster 3, 83%) or entirely (cluster 13, 100%) from AhR-KO tumors. Notably, cluster 13 was the only cluster to express relatively high levels of <u>both</u> granzyme B and perforin (*Prf1*), essential CTL effector molecules. In contrast, *Gzmb* and *Prf1* expression was either not significantly different or significantly lower, relative to all other clusters, in three CD8 clusters predominantly made up of T cells from wildtype tumors, i.e., clusters 2, 6 and 8.

Given that AhR-KO cluster 13 is apparently more active in general and more interactive with APC than T cells from CMT167^WT^ tumors, and that cluster 13 cells express higher *Gzmb* and *Prf1* levels than the aggregate of all the other CD8 clusters, we postulated that it would express a more robust global transcriptomic profile of an active CTL than other CD8 clusters. Gene Set Enrichment Analysis (GSEA) was performed using the clusterProfiler package to test this hypothesis. Three sets of upregulated genes in cluster 13 were generated relative to: 1) all other CD8 clusters, 2) the aggregate of clusters 2, 6, and 8, and 3) cluster 6. Parameters were set to a minimum of 25% expressing cells and a minimum average log2 fold change of 0.25. The FindMarkers function with the MAST wrapper in Seurat was used to rank these DEG sets by decreasing average log2 fold change (avg -log2FC) and adjusted p-value (-log10). The ranked genes were then input into the GSEA enrichment function using the ImmunesigDB gene sets from the MSigDB website. All three sets of genes upregulated in cluster 13 were significantly enriched in six sets of genes experimentally shown to be upregulated in activated CD8 T cells (**Fig. 9F**). These include three sets of genes upregulated in effector CD8 T cells as compared with naïve CD8 T cells (GSE9650 Eff. vs Naïve CD8 T cells, Goldrath Eff. *vs* Naïve CD8 T cells, Kaech Eff. *vs* Naïve CD8 T cells)(81, 82), two sets of genes upregulated in effector CD8 T cells relative to memory CD8 T cells (GSE9650 Eff. *vs* Mem. CD8 T cells, Kaech Eff. *vs* Mem. CD8 T cells)(81, 83) and one set of genes upregulated in activated CD8 T cells as compared with tolerized CD8 T cells (GSE14699 Activated *vs* Tolerant CD8 T)(84)(**Fig. 9F**). All of these results are consistent with the hypothesis that at least some subsets of CD8 T cells from CMT167^AhR-KO^ tumors, including and especially cluster 13, are more active in the tumor microenvironment than CD8 T cells from CMT167^WT^ tumors.

## Discussion

The current studies were motivated in part by our incomplete understanding of factors regulating immune checkpoints, the sometimes contradictory effects of IFNγ on tumor immunity, and the accumulating data indicating an important role for the AhR in immune regulation in the presence or absence of environmental agonists. Recent calls for AhR inhibitors as cancer therapeutics (35, 85–88) and the initiation of cancer clinical trials with AhR inhibitors (89) further add significance to the studies.

Our initial studies of global transcriptomic changes in murine and human LUAD cells pointed to several pathways through which the AhR could control intrinsic drivers of cancer cells as well as regulators of immune cells in the tumor microenvironment (TME). With regard to the former, RNA-seq analysis of AhR-regulated genes revealed the potential for the AhR to control expression of multiple genes implicated in LUAD. For example, *Egfr* expression was reduced 13-16 fold in murine and human LUAD cells, respectively, following AhR knockout (**Fig. 1**). Elevated EGFR activity is a prognostic indicator in LUAD (52) and the EGFR itself is an important therapeutic target (90, 91). Thrombospondin 1 (*Thbs1*), downregulated >45 fold in both CMT167 and A549 cells after AhR knockout, is also a prognostic marker of LUAD outcomes (51, 92). *Colra1*, downregulated 9-219 fold after AhR knockout, can regulate LUAD metastasis (50).

With regard to malignant cell effects on the immune TME, bulk RNA-seq data documented significant downregulation of the immune-related genes *Cccl2*, *Ccl5*, *ITGβ2*, *IDO1*, and *CD274*/*PD-L1* following AhR deletion. The decrease in *Ccl2* after AhR deletion is reminiscent of AhR control of the CCL2 receptor, CCR2, in glioblastoma (93) and with AhR control of CCL2 expression in multiple organs after exposure to the prototypic AhR agonist 2,3,7,8-tetrachlorodibenzo(p)dioxin (TCDD) (94), in non-transformed endothelial cells (95), and in triple negative breast cancer cells (96). AhR-driven expression of two chemokine genes, *Ccl2* and *Ccl5*, could increase recruitment of immunosuppressive monocytes, dendritic cells, and T cell subsets to the TME (97–99). Malignant cell ITGβ2 is associated with suppressive or exhausted T cell infiltration in LUAD (53).

RT-qPCR studies confirmed the bulk RNA-seq data by demonstrating that baseline *Cd274* and *Ido1/2* decreased in AhR-knockout CMT167 cells (**Figs. 3-5**). Conversely, treatment of CMT167 cells with the environmental chemical and AhR agonist B(a)P increased *Cd274* and *Ido1/2* expression (**Fig. 3**). These data expand on previous studies showing that cigarette smoke increases PD-L1 on normal human epithelial cells (35) and help reveal a new mechanism for the well-established immunosuppression generated in animal models by B(a)P and other environmental AhR agonists. Furthermore, the ability of the AhR to regulate baseline levels of PD-L1 and IDO is an important finding with clinical implications. For example, tumors with relatively high AhR levels would be expected to be relatively dependent on the PD-1/PD-L1 axis for immune escape. Indeed, tumors from ∼81% of NSCLC patients who achieve a partial or stable response to Pembrolizumab express relatively high AhR levels whereas tumors from ∼75% of patients that responded poorly or not at all to Pembrolizumab exhibit relatively low AhR levels (35).

Of particular interest were the results with IFNγ that highlighted the likelihood that this cytokine, usually associated with T cell activation and tumor immunity, can also play a negative immunoregulatory role in cancer in general (100, 101) and lung cancer in specific (102). That the effects of IFNγ are mediated at least in part by the AhR is a central and novel finding of these studies. LUAD cells are likely to be exposed to IFNγ *in situ* through infiltrating T cells, NK cells, and/or neutrophils (**Fig. 10**).

**Figure 10.**
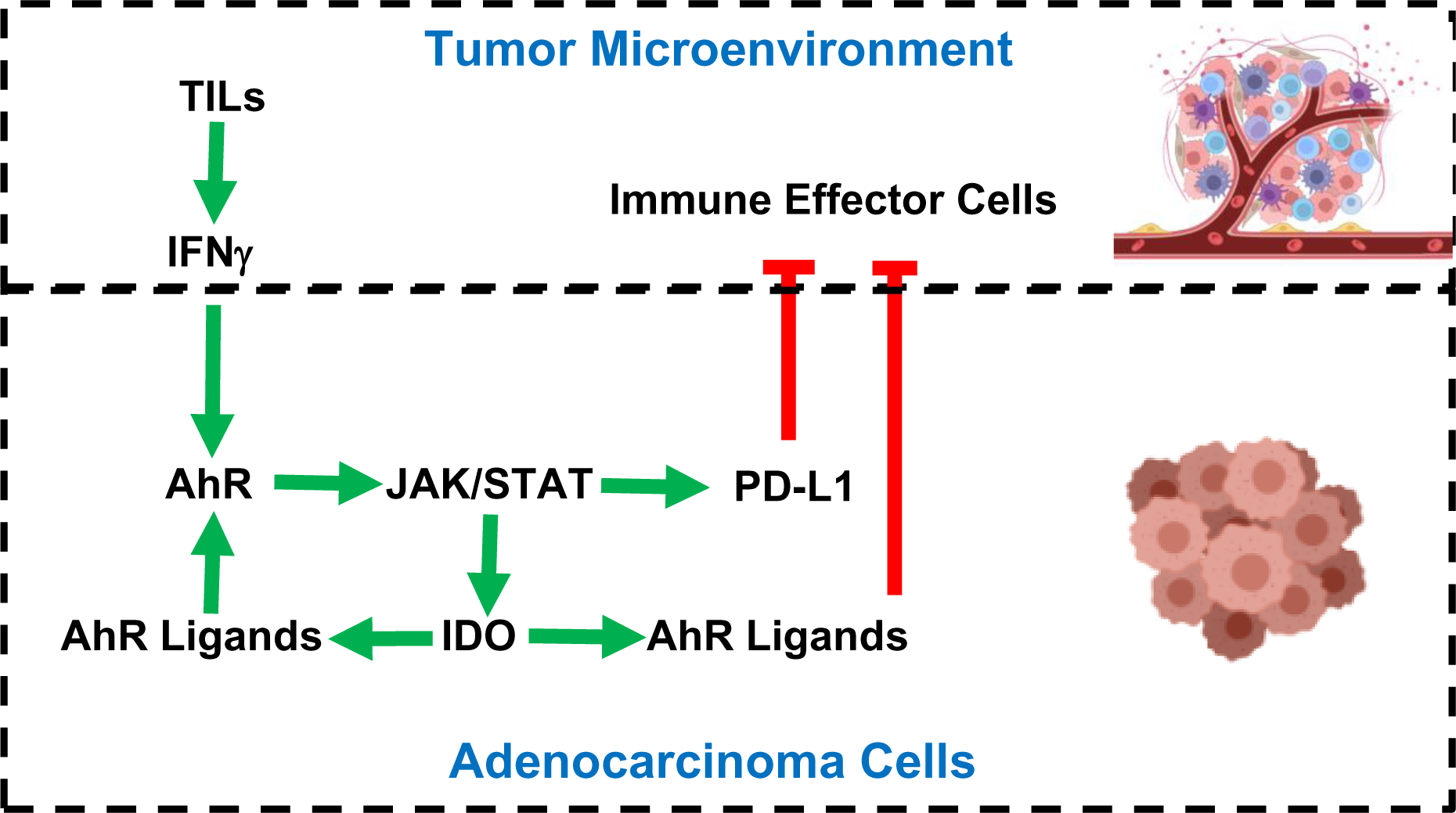
Working model of IFNγ and AhR-mediated regulation of IDO and PD-L1. TILs in the TME generate IFNγ which upregulates AhR activity in malignant cells through an as yet unidentified pathway(s). AhR signaling boosts the JAK/STAT pathway up-regulating IDO and PD-L1. IDO contributes to generation of tryptophan-derived AhR ligands including kynurenine resulting in an AhR amplification loop. Kynurenine and other AhR ligands may activate the AhR in immune cells in the TME skewing them towards immunosuppressive phenotypes. PD-L1 on malignant cells suppresses immune effector cell function.

It is well established that IFNγ upregulates CD274/PD-L1 and IDO through the JAK/STAT pathway (12, 69). Here we demonstrate that optimal expression of JAK/STAT components (JAK2, STAT1, and STAT3) and critical immune checkpoints targeted by JAK/STAT signaling, PD-L1, and IDO, is AhR dependent. These results suggest a novel signaling pathway initiated or exacerbated by environmental or endogenous AhR ligands and leading to suppression of tumor immunity (**Fig. 10**). If confirmed in other cell contexts, the data would also suggest that the AhR may influence a variety of other important JAK/STAT-dependent outcomes not evaluated here.

The data presented here demonstrate that the AhR participates in an amplification loop in LUAD cells by up-regulating IDO1 and IDO2, proximal enzymes in the Kyn pathway of tryptophan metabolism, and resulting in production of endogenous AhR ligands including but probably not limited to Kyn itself (**Fig. 10**). These AhR ligands not only continue to drive AhR activity withing the malignant LUAD cells but may also contribute to immunosuppression in the tumor microenvironment by inducing or recruiting AhR^+^ Treg (103–105), tolerogenic dendritic cells (24, 106), immunosuppressive macrophages (93), or myeloid-derived suppressor cells (107). Thus, in this context, IFNγ can be viewed as a LUAD enabler through AhR regulation.

The finding that IFNγ induced *Cyp1a1* or *Cyp1b1* in CMT167 cells (**Figs. 4,5**) is predicted to reflect IFNγ induction of IDO and the subsequent increase in Kyn (**Figs. 4,5**) or other downstream tryptophan derived AhR ligands. That a cytokine so critical to immunity can induce these canonical AhR-driven genes as well as IDO suggests a novel route to increased CYP1A1 or CYP1B1 metabolic activity in any cell type within an IFNγ-producing leukocyte-infiltrated TME.

One question that remains to be answered is how IFNγ influences AhR activity upstream of AhR-induced IDO up-regulation (**Fig. 10**). There are several, mostly untested possibilities: 1) IFNγ might increase AhR expression levels. However, in our hands, AhR mRNA and protein levels did not increase with addition of IFNγ in either CMT167 or A549 cells (not shown). 2) IFNγ may modify expression or function of AhR-associated proteins including HSP90, p23, Src, AhR interacting protein (AIP), ARNT, or the AhR repressor (AhRR). To our knowledge there is little to no evidence of IFNγ control of p23, Src, AIP, ARNT, or AhRR. Furthermore, in preliminary experiments with IFNγ-stimulated A549 cells we saw no effect of IFNγ on *HSP90*, *p23*, *SRC*, *AIP*, *ARNT*, or *AhRR* mRNAs by RNA-seq. That said, in one report, HSP90 inhibitors blocked IFNγ-induced upregulation of immune checkpoints IDO1 and PD-L1 in pancreatic ductal adenocarcinoma cells (108). 3) IFNγ may affect AhR activity through epigenetic modeling of AhR target genes (109) lowering the threshold to AhR transcriptional activity. 4) In our hands AhR knockout generally does not completely ablate IDO expression (e.g., **Figs. 3,4,5**). Therefore, there may be some AhR- and or JAK/STAT-independent component to IFNγ upregulation of IDO which would be expected to produce AhR ligands that prime the AhR➔IDO➔AhR ligand amplification loop and result in the outcomes studied herein. To that point, STAT1-independent IFNγ signaling has been documented (110). 5) Perhaps the most likely pathway to IFNγ-mediated AhR up-regulation is through NF-κB. IFNγ induces NF-κB signaling (111), components of which can bind to and modify AhR activity (112, 113). Clearly, more experimentation is required to resolve how IFNγ affects AhR signaling.

Given the *in vitro* effects of AhR knockout, it was hypothesized that CMT167^AhR-KO^ cells would either grow more slowly or not all by virtue of an enabled immune system. Indeed, only 23% of the mice grew CMT167^AhR-KO^ tumors (**Fig. 6A**) and those CMT167^AhR-KO^ tumors that did grow grew at a significantly slower pace than control CMT167^Cas9^ or CMT167^WT^ tumors (e.g., **Fig. 6B**). Furthermore, mice re-challenged with CMT167^WT^ cells seven weeks after inoculation with CMT167^AhR-KO^ cells exhibited significant resistance to outgrowth of wildtype tumors (**Fig. 6C,D**) demonstrating some level of immune memory. Again, these results speak to the significance of AhR activity in malignant cells and the effects that this activity has on the immune microenvironment. That some CMT167^AhR-KO^ tumors did escape immune attack suggests that this model could prove useful in cataloguing mechanisms through which LUADs being targeted with immune checkpoint inhibitors escape an otherwise competent immune system. Experiments assessing the mechanism of immune escape are now underway.

Immunofluorescent studies gave the first hint that immune protection against CMT167^AhR-KO^ tumors is likely mediated by CD45^+^ cells. Thus, while nearly absent in CMT167^WT^ tumors, CD45^+^ TILs were plentiful in CMT167^AhR-KO^ tumors (**Fig. 7A**). Immunophenotyping studies confirmed an abundance of CD4^+^ and CD8^+^ T cells in CMT167^AhR-KO^ tumors (**Fig. 7C-E**). Although a significant percentage of these cells expressed PD-1, their continued accumulation in CMT167^AhR-KO^ tumors over time and the single cell characterization of interactive CD8 T cells suggests that they were not exhausted. Rather, they may represent a population of PD1^+^ tumor antigen-specific CTL as seen in LUAD, colorectal cancer, and HNSCC (74, 75).

More granular analysis of the T cell subsets by scRNA-sequencing revealed the presence of 16 clusters of CD3 T cells, a demonstration of the molecular heterogeneity of tumor-infiltrating T cells. Importantly, the distribution of these T cell subsets was significantly different in CMT167^WT^ vs CMT167^AhR-KO^ tumors with one CD8 subset unique to wildtype tumors (cluster 6) and another unique to CMT167^AhR-KO^ tumors (cluster 13). The only cluster that expressed multiple markers of Tregs or exhausted T cells (*Tnfrs9*, *Tgfβ*, *Tigit*, *Pdcd1/Pd-1*, *Ctla4*) was a CD4 cluster composed of predominantly (90%) T cells from wildtype tumors (cluster 5). Similarly, CD8 cluster 6, composed 100% of T cells from wildtype tumors, expressed six markers of exhausted T cells (*Itgav*, *Tnfsrf9*, *Tigit*, *Pdcd-1*, *Lag3*, and *Havcrw**/**Tim3*). In contrast, CD8 cluster 13, composed 100% of T cells from AhR-KO tumors, expressed the highest levels of granzyme B and perforin mRNAs. GSVA analysis indicated that this cluster was more active than CD8 T cells from wildtype tumors and CellChat analyses supported the hypothesis that these cells are actively engaged with APCs in the CMT167^AhR-KO^ microenvironment. Indeed, it is possible that the PD1 expressing cluster 13 cells represents activated tumor-specific CTL (114–116). The only other CD8 subset to express high *Grzb* levels was also predominantly (83%) made up of T cells from CMT167^AhR-KO^ tumors. While additional functional studies would be required to confirm the implied function of these clusters, the data are consistent with the induction of an immunosuppressive TME by CMT^WT/Cas9^ control cells and a more immunocompetent TME in CMT167^AhR-KO^ tumors. We note that these studies do not exclude a role for non-T cells in LUAD AhR-mediated immunosuppression. Analysis of other TIL subsets that are differentially represented in CMT167^WT/Cas9^ and CMT167^AhR-KO^ tumors, including neutrophils, macrophages, dendritic cells, and B cells, is ongoing.

## Acknowledgements

The authors acknowledge the excellent technical assistance provided by Eric Yang.

## Abbreviations

AhR: Aryl Hydrocarbon Receptor
B(a)P: Benzo(a)pyrene
DGE: Differentially gene expression
DEG: Differentially expressed gene(s)
FICZ: 6-formylindolo(3,2-b)carbazole
GSVA: Geneset variation analysis
GSEA: Gene Set Enrichment Analysis
LUAD: Lung adenocarcinoma
Kyn: Kynurenine
NSCLC: Non-small Cell Lung Cancer
TME, scRNA-seq: single cell RNA sequencing
TCDD: 2,3,7,8-tetrachlorodibenzo-p-dioxin Tumor Microenvironment

## Notes

1 Supported by R01ES029136, R01ES033692, and grants from the Find The Cause Breast Cancer Foundation and the Hahnemann Foundation

### Competing Interest Statement

The authors have declared no competing interest.

## References

1. Ferlay, J., H. R. Shin, F. Bray, D. Forman, C. Mathers, and D. M. Parkin. 2010. Estimates of worldwide burden of cancer in 2008: GLOBOCAN 2008. Int J Cancer 127: 2893–2917.

2. Sainz de Aja, J., A. F. M. Dost, and C. F. Kim. 2021. Alveolar progenitor cells and the origin of lung cancer. J Intern Med 289: 629–635.

3. Rivera, G. A., and H. Wakelee. 2016. Lung Cancer in Never Smokers. Adv Exp Med Biol 893: 43–57.

4. Sezer, A., S. Kilickap, M. Gumus, I. Bondarenko, M. Ozguroglu, M. Gogishvili, H. M. Turk, I. Cicin, D. Bentsion, O. Gladkov, P. Clingan, V. Sriuranpong, N. Rizvi, B. Gao, S. Li, S. Lee, K. McGuire, C. I. Chen, T. Makharadze, S. Paydas, M. Nechaeva, F. Seebach, D. M. Weinreich, G. D. Yancopoulos, G. Gullo, I. Lowy, and P. Rietschel. 2021. Cemiplimab monotherapy for first-line treatment of advanced non-small-cell lung cancer with PD-L1 of at least 50%: a multicentre, open-label, global, phase 3, randomised, controlled trial. Lancet 397: 592–604.

5. Paz-Ares, L. G., S. S. Ramalingam, T. E. Ciuleanu, J. S. Lee, L. Urban, R. B. Caro, K. Park, H. Sakai, Y. Ohe, M. Nishio, C. Audigier-Valette, J. A. Burgers, A. Pluzanski, R. Sangha, C. Gallardo, M. Takeda, H. Linardou, L. Lupinacci, K. H. Lee, C. Caserta, M. Provencio, E. Carcereny, G. A. Otterson, M. Schenker, B. Zurawski, A. Alexandru, A. Vergnenegre, J. Raimbourg, K. Feeney, S. W. Kim, H. Borghaei, K. J. O’Byrne, M. D. Hellmann, A. Memaj, F. E. Nathan, J. Bushong, P. Tran, J. R. Brahmer, and M. Reck. 2022. First-Line Nivolumab Plus Ipilimumab in Advanced NSCLC: 4-Year Outcomes From the Randomized, Open-Label, Phase 3 CheckMate 227 Part 1 Trial. J Thorac Oncol 17: 289–308.

6. Opitz, C. A., U. M. Litzenburger, F. Sahm, M. Ott, I. Tritschler, S. Trump, T. Schumacher, L. Jestaedt, D. Schrenk, M. Weller, M. Jugold, G. J. Guillemin, C. L. Miller, C. Lutz, B. Radlwimmer, I. Lehmann, A. von Deimling, W. Wick, and M. Platten. 2011. An endogenous tumour-promoting ligand of the human aryl hydrocarbon receptor. Nature 478: 197–203.

7. Liu, Y., X. Liang, W. Dong, Y. Fang, J. Lv, T. Zhang, R. Fiskesund, J. Xie, J. Liu, X. Yin, X. Jin, D. Chen, K. Tang, J. Ma, H. Zhang, J. Yu, J. Yan, H. Liang, S. Mo, F. Cheng, Y. Zhou, H. Zhang, J. Wang, J. Li, Y. Chen, B. Cui, Z. W. Hu, X. Cao, F. Xiao-Feng Qin, and B. Huang. 2018. Tumor-repopulating cells induce PD-1 expression in CD8(+) T cells by transferring kynurenine and AhR activation. Cancer Cell 33: 480–494 e487.

8. Amobi-McCloud, A., R. Muthuswamy, S. Battaglia, H. Yu, T. Liu, J. Wang, V. Putluri, P. K. Singh, F. Qian, R. Y. Huang, N. Putluri, T. Tsuji, A. A. Lugade, S. Liu, and K. Odunsi. 2021. IDO1 Expression in Ovarian Cancer Induces PD-1 in T Cells via Aryl Hydrocarbon Receptor Activation. Front Immunol 12: 678999.

9. Litzenburger, U. M., C. A. Opitz, F. Sahm, K. J. Rauschenbach, S. Trump, M. Winter, M. Ott, K. Ochs, C. Lutz, X. Liu, N. Anastasov, I. Lehmann, T. Hofer, A. von Deimling, W. Wick, and M. Platten. 2014. Constitutive IDO expression in human cancer is sustained by an autocrine signaling loop involving IL-6, STAT3 and the AHR. Oncotarget 5: 1038–1051.

10. Mishra, A. K., T. Kadoishi, X. Wang, E. Driver, Z. Chen, X. J. Wang, and J. H. Wang. 2016. Squamous cell carcinomas escape immune surveillance via inducing chronic activation and exhaustion of CD8+ T Cells co-expressing PD-1 and LAG-3 inhibitory receptors. Oncotarget 7: 81341–81356.

11. Zhang, X., X. Liu, W. Zhou, Q. Du, M. Yang, Y. Ding, and R. Hu. 2021. Blockade of IDO-Kynurenine-AhR Axis Ameliorated Colitis-Associated Colon Cancer via Inhibiting Immune Tolerance. Cell Mol Gastroenterol Hepatol 12: 1179–1199.

12. Mandai, M., J. Hamanishi, K. Abiko, N. Matsumura, T. Baba, and I. Konishi. 2016. Dual Faces of IFNgamma in Cancer Progression: A Role of PD-L1 Induction in the Determination of Pro- and Antitumor Immunity. Clin Cancer Res 22: 2329–2334.

13. Mimura, K., J. L. Teh, H. Okayama, K. Shiraishi, L. F. Kua, V. Koh, D. T. Smoot, H. Ashktorab, T. Oike, Y. Suzuki, Z. Fazreen, B. R. Asuncion, A. Shabbir, W. P. Yong, J. So, R. Soong, and K. Kono. 2018. PD-L1 expression is mainly regulated by interferon gamma associated with JAK-STAT pathway in gastric cancer. Cancer Sci 109: 43–53.

14. Gu, Y. Z., J. B. Hogenesch, and C. A. Bradfield. 2000. The PAS superfamily: sensors of environmental and developmental signals. Annu Rev Pharmacol Toxicol 40: 519–561.

15. Holme, J. A., J. Vondracek, M. Machala, D. Lagadic-Gossmann, C. F. A. Vogel, E. Le Ferrec, L. Sparfel, and J. Ovrevik. 2023. Lung cancer associated with combustion particles and fine particulate matter (PM(2.5)) - The roles of polycyclic aromatic hydrocarbons (PAHs) and the aryl hydrocarbon receptor (AhR). Biochem Pharmacol 216: 115801.

16. Tsay, J. J., K. M. Tchou-Wong, A. K. Greenberg, H. Pass, and W. N. Rom. 2013. Aryl hydrocarbon receptor and lung cancer. Anticancer Res 33: 1247–1256.

17. Mandal, P. K. 2005. Dioxin: a review of its environmental effects and its aryl hydrocarbon receptor biology. J Comp Physiol B 175: 221–230.

18. Nebert, D. W., A. Puga, and V. Vasiliou. 1993. Role of the Ah receptor and the dioxin-inducible [Ah] gene battery in toxicity, cancer, and signal transduction. Ann N Y Acad Sci 685: 624–640.

19. Quintana, F. J. 2013. Regulation of central nervous system autoimmunity by the aryl hydrocarbon receptor. Semin Immunopathol 35: 627–635.

20. Bruhs, A., T. Haarmann-Stemmann, K. Frauenstein, J. Krutmann, T. Schwarz, and A. Schwarz. 2015. Activation of the arylhydrocarbon receptor causes immunosuppression primarily by modulating dendritic cells. J Invest Dermatol 135: 435–444.

21. Campesato, L. F., S. Budhu, J. Tchaicha, C. H. Weng, M. Gigoux, I. J. Cohen, D. Redmond, L. Mangarin, S. Pourpe, C. Liu, R. Zappasodi, D. Zamarin, J. Cavanaugh, A. C. Castro, M. G. Manfredi, K. McGovern, T. Merghoub, and J. D. Wolchok. 2020. Blockade of the AHR restricts a Treg-macrophage suppressive axis induced by L-Kynurenine. Nat Commun 11: 4011.

22. Mezrich, J. D., J. H. Fechner, X. Zhang, B. P. Johnson, W. J. Burlingham, and C. A. Bradfield. 2010. An interaction between kynurenine and the aryl hydrocarbon receptor can generate regulatory T cells. J Immunol 185: 3190–3198.

23. Quintana, F. J., and D. H. Sherr. 2013. Aryl hydrocarbon receptor control of adaptive immunity. Pharmacological reviews 65: 1148–1161.

24. Nguyen, N. T., A. Kimura, T. Nakahama, I. Chinen, K. Masuda, K. Nohara, Y. Fujii-Kuriyama, and T. Kishimoto. 2010. Aryl hydrocarbon receptor negatively regulates dendritic cell immunogenicity via a kynurenine-dependent mechanism. Proc Natl Acad Sci U S A 107: 19961–19966.

25. Moyer, B. J., I. Y. Rojas, I. A. Murray, S. Lee, H. F. Hazlett, G. H. Perdew, and C. R. Tomlinson. 2017. Indoleamine 2,3-dioxygenase 1 (IDO1) inhibitors activate the aryl hydrocarbon receptor. Toxicol Appl Pharmacol 323: 74–80.

26. Murray, I. A., A. D. Patterson, and G. H. Perdew. 2014. Aryl hydrocarbon receptor ligands in cancer: friend and foe. Nat Rev Cancer 14: 801–814.

27. Novikov, O., Z. Wang, E. A. Stanford, A. J. Parks, A. Ramirez-Cardenas, E. Landesman, I. Laklouk, C. Sarita-Reyes, D. Gusenleitner, A. Li, S. Monti, S. Manteiga, K. Lee, and D. H. Sherr. 2016. An aryl hydrocarbon receptor-mediated amplification loop that enforces cell migration in ER-/PR-/Her2-human breast cancer cells. Mol Pharmacol 90: 674–688.

28. Stanford, E. A., Z. Wang, O. Novikov, F. Mulas, E. Landesman-Bollag, S. Monti, B. W. Smith, D. C. Seldin, G. J. Murphy, and D. H. Sherr. 2016. The role of the aryl hydrocarbon receptor in the development of cells with the molecular and functional characteristics of cancer stem-like cells. BMC Biol 14: 20.

29. Prud’homme, G. J. 2012. Cancer stem cells and novel targets for antitumor strategies. Curr Pharm Des 18: 2838–2849.

30. Therachiyil, L., R. Krishnankutty, F. Ahmad, J. M. Mateo, S. Uddin, and H. M. Korashy. 2022. Aryl Hydrocarbon Receptor Promotes Cell Growth, Stemness Like Characteristics, and Metastasis in Human Ovarian Cancer via Activation of PI3K/Akt, beta-Catenin, and Epithelial to Mesenchymal Transition Pathways. Int J Mol Sci 23.

31. Rejano-Gordillo, C., A. Ordiales-Talavero, A. Nacarino-Palma, J. M. Merino, F. J. Gonzalez-Rico, and P. M. Fernandez-Salguero. 2022. Aryl Hydrocarbon Receptor: From Homeostasis to Tumor Progression. Front Cell Dev Biol 10: 884004.

32. Ye, M., Y. Zhang, H. Gao, Y. Xu, P. Jing, J. Wu, X. Zhang, J. Xiong, C. Dong, L. Yao, J. Zhang, and J. Zhang. 2018. Activation of the Aryl Hydrocarbon Receptor Leads to Resistance to EGFR TKIs in Non-Small Cell Lung Cancer by Activating Src-mediated Bypass Signaling. Clin Cancer Res 24: 1227–1239.

33. Vrzalova, A., P. Pecinkova, P. Illes, S. Gurska, P. Dzubak, M. Szotkowski, M. Hajduch, S. Mani, and Z. Dvorak. 2022. Mixture Effects of Tryptophan Intestinal Microbial Metabolites on Aryl Hydrocarbon Receptor Activity. Int J Mol Sci 23.

34. Hubbard, T. D., I. A. Murray, W. H. Bisson, T. S. Lahoti, K. Gowda, S. G. Amin, A. D. Patterson, and G. H. Perdew. 2015. Adaptation of the human aryl hydrocarbon receptor to sense microbiota-derived indoles. Sci Rep 5: 12689.

35. Wang, G. Z., L. Zhang, X. C. Zhao, S. H. Gao, L. W. Qu, H. Yu, W. F. Fang, Y. C. Zhou, F. Liang, C. Zhang, Y. C. Huang, Z. Liu, Y. X. Fu, and G. B. Zhou. 2019. The aryl hydrocarbon receptor mediates tobacco-induced PD-L1 expression and is associated with response to immunotherapy. Nat Commun 10: 1125.

36. Kenison, J. E., Z. Wang, K. Yang, M. Snyder, F. J. Quintana, and D. H. Sherr. 2021. The aryl hydrocarbon receptor suppresses immunity to oral squamous cell carcinoma through immune checkpoint regulation. Proc Natl Acad Sci U S A 118.

37. Li, H. Y., M. McSharry, B. Bullock, T. T. Nguyen, J. Kwak, J. M. Poczobutt, T. R. Sippel, L. E. Heasley, M. C. Weiser-Evans, E. T. Clambey, and R. A. Nemenoff. 2017. The Tumor Microenvironment Regulates Sensitivity of Murine Lung Tumors to PD-1/PD-L1 Antibody Blockade. Cancer Immunol Res 5: 767–777.

38. Liao, H., X. Chang, L. Gao, C. Ye, Y. Qiao, L. Xie, J. Lin, S. Cai, and H. Dong. 2023. IL-17A promotes tumorigenesis and upregulates PD-L1 expression in non-small cell lung cancer. J Transl Med 21: 828.

39. Nawas, A. F., A. Solmonson, B. Gao, R. J. DeBerardinis, J. D. Minna, M. Conacci-Sorrell, and C. R. Mendelson. 2023. IL-1beta mediates the induction of immune checkpoint regulators IDO1 and PD-L1 in lung adenocarcinoma cells. Cell Commun Signal 21: 331.

40. Sanjana, N. E., O. Shalem, and F. Zhang. 2014. Improved vectors and genome-wide libraries for CRISPR screening. Nat Methods 11: 783–784.

41. Hong, R., Y. Koga, S. Bandyadka, A. Leshchyk, Y. Wang, V. Akavoor, X. Cao, I. Sarfraz, Z. Wang, S. Alabdullatif, F. Jansen, M. Yajima, W. E. Johnson, and J. D. Campbell. 2022. Comprehensive generation, visualization, and reporting of quality control metrics for single-cell RNA sequencing data. Nat Commun 13: 1688.

42. Aran, D., A. P. Looney, L. Liu, E. Wu, V. Fong, A. Hsu, S. Chak, R. P. Naikawadi, P. J. Wolters, A. R. Abate, A. J. Butte, and M. Bhattacharya. 2019. Reference-based analysis of lung single-cell sequencing reveals a transitional profibrotic macrophage. Nat Immunol 20: 163–172.

43. Heng, T. S., M. W. Painter, and C. Immunological Genome Project. 2008. The Immunological Genome Project: networks of gene expression in immune cells. Nat Immunol 9: 1091–1094.

44. Stuart, T., A. Butler, P. Hoffman, C. Hafemeister, E. Papalexi, W. M. Mauck, 3rd, Y. Hao, M. Stoeckius, P. Smibert, and R. Satija. 2019. Comprehensive Integration of Single-Cell Data. Cell 177: 1888–1902 e1821.

45. Jin, S., C. F. Guerrero-Juarez, L. Zhang, I. Chang, R. Ramos, C. H. Kuan, P. Myung, M. V. Plikus, and Q. Nie. 2021. Inference and analysis of cell-cell communication using CellChat. Nat Commun 12: 1088.

46. Hanzelmann, S., R. Castelo, and J. Guinney. 2013. GSVA: gene set variation analysis for microarray and RNA-seq data. BMC Bioinformatics 14: 7.

47. Newman, A. M., C. B. Steen, C. L. Liu, A. J. Gentles, A. A. Chaudhuri, F. Scherer, M. S. Khodadoust, M. S. Esfahani, B. A. Luca, D. Steiner, M. Diehn, and A. A. Alizadeh. 2019. Determining cell type abundance and expression from bulk tissues with digital cytometry. Nat Biotechnol 37: 773–782.

48. Finak, G., A. McDavid, M. Yajima, J. Deng, V. Gersuk, A. K. Shalek, C. K. Slichter, H. W. Miller, M. J. McElrath, M. Prlic, P. S. Linsley, and R. Gottardo. 2015. MAST: a flexible statistical framework for assessing transcriptional changes and characterizing heterogeneity in single-cell RNA sequencing data. Genome Biol 16: 278.

49. Huang, X., Q. Sun, C. Chen, Y. Zhang, X. Kang, J. Y. Zhang, D. W. Ma, L. Xia, L. Xu, X. Y. Xu, and B. H. Ren. 2017. MUC1 overexpression predicts worse survival in patients with non-small cell lung cancer: evidence from an updated meta-analysis. Oncotarget 8: 90315–90326.

50. Liu, W., H. Wei, Z. Gao, G. Chen, Y. Liu, X. Gao, G. Bai, S. He, T. Liu, W. Xu, X. Yang, J. Jiao, and J. Xiao. 2018. COL5A1 may contribute the metastasis of lung adenocarcinoma. Gene 665: 57–66.

51. Sun, R., X. Meng, W. Wang, B. Liu, X. Lv, J. Yuan, L. Zeng, Y. Chen, B. Yuan, and S. Yang. 2019. Five genes may predict metastasis in non-small cell lung cancer using bioinformatics analysis. Oncol Lett 18: 1723–1732.

52. Gao, M. G., S. Z. Wang, K. H. Han, S. N. Xie, and Q. Y. Liu. 2022. Clinical characteristics and prognostic value of EGFR mutation in stage I lung adenocarcinoma with spread through air spaces after surgical resection. Neoplasma 69: 1480–1489.

53. Wu, J., W. Wang, Z. Li, and X. Ye. 2022. The prognostic and immune infiltration role of ITGB superfamily members in non-small cell lung cancer. Am J Transl Res 14: 6445–6466.

54. Zu, L., J. He, N. Zhou, J. Zeng, Y. Zhu, Q. Tang, X. Jin, L. Zhang, and S. Xu. 2022. The Profile and Clinical Significance of ITGB2 Expression in Non-Small-Cell Lung Cancer. J Clin Med 11.

55. Lee, K. Y., P. W. Shueng, C. M. Chou, B. X. Lin, M. H. Lin, D. Y. Kuo, I. L. Tsai, S. M. Wu, and C. W. Lin. 2020. Elevation of CD109 promotes metastasis and drug resistance in lung cancer via activation of EGFR-AKT-mTOR signaling. Cancer Sci 111: 1652–1662.

56. Fridlender, Z. G., V. Kapoor, G. Buchlis, G. Cheng, J. Sun, L. C. Wang, S. Singhal, L. A. Snyder, and S. M. Albelda. 2011. Monocyte chemoattractant protein-1 blockade inhibits lung cancer tumor growth by altering macrophage phenotype and activating CD8+ cells. Am J Respir Cell Mol Biol 44: 230–237.

57. Melese, E. S., E. Franks, R. A. Cederberg, B. T. Harbourne, R. Shi, B. J. Wadsworth, J. L. Collier, E. C. Halvorsen, F. Johnson, J. Luu, M. H. Oh, V. Lam, G. Krystal, S. B. Hoover, M. Raffeld, R. M. Simpson, A. M. Unni, W. L. Lam, S. Lam, N. Abraham, K. L. Bennewith, and W. W. Lockwood. 2022. CCL5 production in lung cancer cells leads to an altered immune microenvironment and promotes tumor development. Oncoimmunology 11: 2010905.

58. Kocher, F., A. Amann, K. Zimmer, S. Geisler, D. Fuchs, R. Pichler, D. Wolf, K. Kurz, A. Seeber, and A. Pircher. 2021. High indoleamine-2,3-dioxygenase 1 (IDO) activity is linked to primary resistance to immunotherapy in non-small cell lung cancer (NSCLC). Transl Lung Cancer Res 10: 304–313.

59. Han, Y., Y. Zhang, Y. Tian, M. Zhang, C. Xiang, Q. Zhen, J. Liu, Y. Shang, Y. Zhao, H. Si, and A. Sui. 2022. The Interaction of the IFNgamma/JAK/STAT1 and JAK/STAT3 Signalling Pathways in EGFR-Mutated Lung Adenocarcinoma Cells. J Oncol 2022: 9016296.

60. Kado, S. Y., K. Bein, A. R. Castaneda, A. A. Pouraryan, N. Garrity, Y. Ishihara, A. Rossi, T. Haarmann-Stemmann, C. A. Sweeney, and C. F. A. Vogel. 2023. Regulation of IDO2 by the Aryl Hydrocarbon Receptor (AhR) in Breast Cancer. Cells 12.

61. Bekki, K., H. Vogel, W. Li, T. Ito, C. Sweeney, T. Haarmann-Stemmann, F. Matsumura, and C. F. Vogel. 2015. The aryl hydrocarbon receptor (AhR) mediates resistance to apoptosis induced in breast cancer cells. Pestic Biochem Physiol 120: 5–13.

62. Vogel, C. F., S. R. Goth, B. Dong, I. N. Pessah, and F. Matsumura. 2008. Aryl hydrocarbon receptor signaling mediates expression of indoleamine 2,3-dioxygenase. Biochem Biophys Res Commun 375: 331–335.

63. Bankoti, J., B. Rase, T. Simones, and D. M. Shepherd. 2010. Functional and phenotypic effects of AhR activation in inflammatory dendritic cells. Toxicol Appl Pharmacol 246: 18–28.

64. Muller, A. J., M. G. Manfredi, Y. Zakharia, and G. C. Prendergast. 2019. Inhibiting IDO pathways to treat cancer: lessons from the ECHO-301 trial and beyond. Semin Immunopathol 41: 41–48.

65. Fujiwara, Y., S. Kato, M. K. Nesline, J. M. Conroy, P. DePietro, S. Pabla, and R. Kurzrock. 2022. Indoleamine 2,3-dioxygenase (IDO) inhibitors and cancer immunotherapy. Cancer Treat Rev 110: 102461.

66. Powderly, J. D., S. J. Klempner, A. Naing, J. Bendell, I. Garrido-Laguna, D. V. T. Catenacci, M. H. Taylor, J. J. Lee, F. Zheng, F. Zhou, X. Gong, H. Gowda, and G. L. Beatty. 2022. Epacadostat Plus Pembrolizumab and Chemotherapy for Advanced Solid Tumors: Results from the Phase I/II ECHO-207/KEYNOTE-723 Study. Oncologist 27: 905–e848.

67. Zhang, Y., Z. Hu, J. Zhang, C. Ren, and Y. Wang. 2022. Dual-target inhibitors of indoleamine 2, 3 dioxygenase 1 (Ido1): A promising direction in cancer immunotherapy. Eur J Med Chem 238: 114524.

68. El Jamal, S. M., E. B. Taylor, Z. Y. Abd Elmageed, A. A. Alamodi, D. Selimovic, A. Alkhateeb, M. Hannig, S. Y. Hassan, S. Santourlidis, P. L. Friedlander, Y. Haikel, S. Vijaykumar, E. Kandil, and M. Hassan. 2016. Interferon gamma-induced apoptosis of head and neck squamous cell carcinoma is connected to indoleamine-2,3-dioxygenase via mitochondrial and ER stress-associated pathways. Cell Div 11: 11.

69. Zhu, J., Y. Li, and X. Lv. 2022. IL4I1 enhances PD-L1 expression through JAK/STAT signaling pathway in lung adenocarcinoma. Immunogenetics.

70. Iwasaki, T., K. Kohashi, Y. Toda, S. Ishihara, Y. Yamada, and Y. Oda. 2021. Association of PD-L1 and IDO1 expression with JAK-STAT pathway activation in soft-tissue leiomyosarcoma. J Cancer Res Clin Oncol 147: 1451–1463.

71. Kazan, O., G. Kir, M. Culpan, G. E. Cecikoglu, G. Atis, and A. Yildirim. 2022. The association between PI3K, JAK/STAT pathways with the PDL-1 expression in prostate cancer. Andrologia 54: e14541.

72. Li, C., F. Yang, R. Wang, W. Li, N. Maskey, W. Zhang, Y. Guo, S. Liu, H. Wang, and X. Yao. 2021. CALD1 promotes the expression of PD-L1 in bladder cancer via the JAK/STAT signaling pathway. Ann Transl Med 9: 1441.

73. Horvath, C. M. 2004. The Jak-STAT pathway stimulated by interferon gamma. Sci STKE 2004: tr8.

74. Thommen, D. S., V. H. Koelzer, P. Herzig, A. Roller, M. Trefny, S. Dimeloe, A. Kiialainen, J. Hanhart, C. Schill, C. Hess, S. Savic Prince, M. Wiese, D. Lardinois, P. C. Ho, C. Klein, V. Karanikas, K. D. Mertz, T. N. Schumacher, and A. Zippelius. 2018. A transcriptionally and functionally distinct PD-1(+) CD8(+) T cell pool with predictive potential in non-small-cell lung cancer treated with PD-1 blockade. Nat Med 24: 994–1004.

75. Duhen, R., O. Fesneau, K. A. Samson, A. K. Frye, M. Beymer, V. Rajamanickam, D. Ross, E. Tran, B. Bernard, A. D. Weinberg, and T. Duhen. 2022. PD-1 and ICOS coexpression identifies tumor-reactive CD4+ T cells in human solid tumors. J Clin Invest 132.

76. Alspach, E., D. M. Lussier, A. P. Miceli, I. Kizhvatov, M. DuPage, A. M. Luoma, W. Meng, C. F. Lichti, E. Esaulova, A. N. Vomund, D. Runci, J. P. Ward, M. M. Gubin, R. F. V. Medrano, C. D. Arthur, J. M. White, K. C. F. Sheehan, A. Chen, K. W. Wucherpfennig, T. Jacks, E. R. Unanue, M. N. Artyomov, and R. D. Schreiber. 2019. MHC-II neoantigens shape tumour immunity and response to immunotherapy. Nature 574: 696–701.

77. Cho, J. W., J. Son, S. J. Ha, and I. Lee. 2021. Systems biology analysis identifies TNFRSF9 as a functional marker of tumor-infiltrating regulatory T-cell enabling clinical outcome prediction in lung cancer. Comput Struct Biotechnol J 19: 860–868.

78. Miragaia, R. J., T. Gomes, A. Chomka, L. Jardine, A. Riedel, A. N. Hegazy, N. Whibley, A. Tucci, X. Chen, I. Lindeman, G. Emerton, T. Krausgruber, J. Shields, M. Haniffa, F. Powrie, and S. A. Teichmann. 2019. Single-Cell Transcriptomics of Regulatory T Cells Reveals Trajectories of Tissue Adaptation. Immunity 50: 493–504 e497.

79. Mair, I., S. E. J. Zandee, I. S. Toor, L. Saul, R. C. McPherson, M. D. Leech, D. J. Smyth, R. A. O’Connor, N. C. Henderson, and S. M. Anderton. 2018. A Context-Dependent Role for alphav Integrins in Regulatory T Cell Accumulation at Sites of Inflammation. Front Immunol 9: 264.

80. Edwards, J. P., A. M. Thornton, and E. M. Shevach. 2014. Release of active TGF-beta1 from the latent TGF-beta1/GARP complex on T regulatory cells is mediated by integrin beta8. J Immunol 193: 2843–2849.

81. Kaech, S. M., S. Hemby, E. Kersh, and R. Ahmed. 2002. Molecular and functional profiling of memory CD8 T cell differentiation. Cell 111: 837–851.

82. Luckey, C. J., D. Bhattacharya, A. W. Goldrath, I. L. Weissman, C. Benoist, and D. Mathis. 2006. Memory T and memory B cells share a transcriptional program of self-renewal with long-term hematopoietic stem cells. Proc Natl Acad Sci U S A 103: 3304–3309.

83. Wherry, E. J., S. J. Ha, S. M. Kaech, W. N. Haining, S. Sarkar, V. Kalia, S. Subramaniam, J. N. Blattman, D. L. Barber, and R. Ahmed. 2007. Molecular signature of CD8+ T cell exhaustion during chronic viral infection. Immunity 27: 670–684.

84. Parish, I. A., S. Rao, G. K. Smyth, T. Juelich, G. S. Denyer, G. M. Davey, A. Strasser, and W. R. Heath. 2009. The molecular signature of CD8+ T cells undergoing deletional tolerance. Blood 113: 4575–4585.

85. Paris, A., N. Tardif, F. M. Baietti, C. Berra, H. M. Leclair, E. Leucci, M. D. Galibert, and S. Corre. 2022. The AhR-SRC axis as a therapeutic vulnerability in BRAFi-resistant melanoma. EMBO Mol Med 14: e15677.

86. Morrow, D., C. Qin, R. Smith, 3rd, and S. Safe. 2004. Aryl hydrocarbon receptor-mediated inhibition of LNCaP prostate cancer cell growth and hormone-induced transactivation. J Steroid Biochem Mol Biol 88: 27–36.

87. Martano, M., P. Stiuso, A. Facchiano, S. De Maria, D. Vanacore, B. Restucci, C. Rubini, M. Caraglia, G. Ravagnan, and L. Lo Muzio. 2018. Aryl hydrocarbon receptor, a tumor grade associated marker of oral cancer, is directly downregulated by polydatin: A pilot study. Oncol Rep 40: 1435–1442.

88. McGovern, K., A. C. Castro, J. Cavanaugh, S. Coma, M. Walsh, J. Tchaicha, S. Syed, P. Natarajan, M. Manfredi, X. M. Zhang, and J. Ecsedy. 2022. Discovery and Characterization of a Novel Aryl Hydrocarbon Receptor Inhibitor, IK-175, and Its Inhibitory Activity on Tumor Immune Suppression. Mol Cancer Ther 21: 1261–1272.

89. McKean, M., D. Aggen, N. Lakhani, B. Bashir, J. Luke, J. Hoffman-Censits, D. Alhalabi, I. Bowman, E. Guancial, A. Tan, T. Lingaraj, M. Timothy, K. Kacena, K. Malek, and S. Santillana. 2022. Phase 1a/b open-label study of IK-175, an oral AHR inhibitor, alone and in combination with nivolumab in patients with locally advanced or metastatic solid tumors and urothelial carcinoma. J. Clin. Onc. 40.

90. Li, J., P. Chen, Q. Wu, L. Guo, K. W. Leong, K. I. Chan, and H. F. Kwok. 2022. A novel combination treatment of antiADAM17 antibody and erlotinib to overcome acquired drug resistance in non-small cell lung cancer through the FOXO3a/FOXM1 axis. Cell Mol Life Sci 79: 614.

91. Lee, Y., H. R. Kim, M. H. Hong, K. H. Lee, K. U. Park, G. K. Lee, H. Y. Kim, S. H. Lee, K. Y. Lim, S. J. Yoon, B. C. Cho, and J. Y. Han. 2023. A randomized Phase 2 study to compare erlotinib with or without bevacizumab in previously untreated patients with advanced non-small cell lung cancer with EGFR mutation. Cancer 129: 405–414.

92. Li, R., X. Liu, X. J. Zhou, X. Chen, J. P. Li, Y. H. Yin, and Y. Q. Qu. 2020. Identification and validation of the prognostic value of immune-related genes in non-small cell lung cancer. Am J Transl Res 12: 5844–5865.

93. Takenaka, M. C., G. Gabriely, V. Rothhammer, I. D. Mascanfroni, M. A. Wheeler, C. C. Chao, C. Gutierrez-Vazquez, J. Kenison, E. C. Tjon, A. Barroso, T. Vandeventer, K. A. de Lima, S. Rothweiler, L. Mayo, S. Ghannam, S. Zandee, L. Healy, D. Sherr, M. F. Farez, A. Prat, J. Antel, D. A. Reardon, H. Zhang, S. C. Robson, G. Getz, H. L. Weiner, and F. J. Quintana. 2020. Control of tumor-associated macrophages and T cells in glioblastoma via AHR and CD39. Nat Neurosci 22: 729–740.

94. Vogel, C. F., N. Nishimura, E. Sciullo, P. Wong, W. Li, and F. Matsumura. 2007. Modulation of the chemokines KC and MCP-1 by 2,3,7,8-tetrachlorodibenzo-p-dioxin (TCDD) in mice. Arch Biochem Biophys 461: 169–175.

95. Watanabe, I., J. Tatebe, S. Namba, M. Koizumi, J. Yamazaki, and T. Morita. 2013. Activation of aryl hydrocarbon receptor mediates indoxyl sulfate-induced monocyte chemoattractant protein-1 expression in human umbilical vein endothelial cells. Circ J 77: 224–230.

96. Goode, G., S. Pratap, and S. E. Eltom. 2014. Depletion of the aryl hydrocarbon receptor in MDA-MB-231 human breast cancer cells altered the expression of genes in key regulatory pathways of cancer. PLoS One 9: e100103.

97. Li, L., Y. D. Liu, Y. T. Zhan, Y. H. Zhu, Y. Li, D. Xie, and X. Y. Guan. 2018. High levels of CCL2 or CCL4 in the tumor microenvironment predict unfavorable survival in lung adenocarcinoma. Thorac Cancer 9: 775–784.

98. Larroquette, M., J. P. Guegan, B. Besse, S. Cousin, M. Brunet, S. Le Moulec, F. Le Loarer, C. Rey, J. C. Soria, F. Barlesi, A. Bessede, J. Y. Scoazec, I. Soubeyran, and A. Italiano. 2022. Spatial transcriptomics of macrophage infiltration in non-small cell lung cancer reveals determinants of sensitivity and resistance to anti-PD1/PD-L1 antibodies. J Immunother Cancer 10.

99. Zhang, M., W. Yang, P. Wang, Y. Deng, Y. T. Dong, F. F. Liu, R. Huang, P. Zhang, Y. Q. Duan, X. D. Liu, D. Lin, Q. Chu, and B. Zhong. 2020. CCL7 recruits cDC1 to promote antitumor immunity and facilitate checkpoint immunotherapy to non-small cell lung cancer. Nat Commun 11: 6119.

100. Benci, J. L., L. R. Johnson, R. Choa, Y. Xu, J. Qiu, Z. Zhou, B. Xu, D. Ye, K. L. Nathanson, C. H. June, E. J. Wherry, N. R. Zhang, H. Ishwaran, M. D. Hellmann, J. D. Wolchok, T. Kambayashi, and A. J. Minn. 2019. Opposing Functions of Interferon Coordinate Adaptive and Innate Immune Responses to Cancer Immune Checkpoint Blockade. Cell 178: 933–948 e914.

101. Jing, Z. L., G. L. Liu, N. Zhou, D. Y. Xu, N. Feng, Y. Lei, L. L. Ma, M. S. Tang, G. H. Tong, N. Tang, and Y. J. Deng. 2024. Interferon-gamma in the tumor microenvironment promotes the expression of B7H4 in colorectal cancer cells, thereby inhibiting cytotoxic T cells. Sci Rep 14: 6053.

102. Zagorulya, M., L. Yim, D. M. Morgan, A. Edwards, E. Torres-Mejia, N. Momin, C. V. McCreery, I. L. Zamora, B. L. Horton, J. G. Fox, K. D. Wittrup, J. C. Love, and S. Spranger. 2023. Tissue-specific abundance of interferon-gamma drives regulatory T cells to restrain DC1-mediated priming of cytotoxic T cells against lung cancer. Immunity 56: 386–405 e310.

103. Quintana, F. J., A. S. Basso, A. H. Iglesias, T. Korn, M. F. Farez, E. Bettelli, M. Caccamo, M. Oukka, and H. L. Weiner. 2008. Control of T(reg) and T(H)17 cell differentiation by the aryl hydrocarbon receptor. Nature 453: 65–71.

104. Abron, J. D., N. P. Singh, M. K. Mishra, R. L. Price, M. Nagarkatti, P. S. Nagarkatti, and U. P. Singh. 2018. An endogenous aryl hydrocarbon receptor (AhR) ligand, ITE induces regulatory T cells (Tregs) and ameliorates experimental colitis. Am J Physiol Gastrointest Liver Physiol.

105. Apetoh, L., F. J. Quintana, C. Pot, N. Joller, S. Xiao, D. Kumar, E. J. Burns, D. H. Sherr, H. L. Weiner, and V. K. Kuchroo. 2010. The aryl hydrocarbon receptor interacts with c-Maf to promote the differentiation of type 1 regulatory T cells induced by IL-27. Nat Immunol 11: 854–861.

106. Quintana, F. J., G. Murugaiyan, M. F. Farez, M. Mitsdoerffer, A. M. Tukpah, E. J. Burns, and H. L. Weiner. 2010. An endogenous aryl hydrocarbon receptor ligand acts on dendritic cells and T cells to suppress experimental autoimmune encephalomyelitis. Proc Natl Acad Sci U S A.

107. Neamah, W. H., N. P. Singh, H. Alghetaa, O. A. Abdulla, S. Chatterjee, P. B. Busbee, M. Nagarkatti, and P. Nagarkatti. 2019. AhR activation leads to massive mobilization of myeloid-derived suppressor cells with immunosuppressive activity through regulation of CXCR2 and microRNA miR-150-5p and miR-543-3p that target anti-inflammatory genes. J Immunol 203: 1830–1844.

108. Liu, K., J. Huang, J. Liu, C. Li, G. Kroemer, D. Tang, and R. Kang. 2022. HSP90 Mediates IFNgamma-Induced Adaptive Resistance to Anti-PD-1 Immunotherapy. Cancer Res 82: 2003–2018.

109. Ivashkiv, L. B. 2018. IFNgamma: signalling, epigenetics and roles in immunity, metabolism, disease and cancer immunotherapy. Nat Rev Immunol 18: 545–558.

110. Ramana, C. V., M. P. Gil, Y. Han, R. M. Ransohoff, R. D. Schreiber, and G. R. Stark. 2001. Stat1-independent regulation of gene expression in response to IFN-gamma. Proc Natl Acad Sci U S A 98: 6674–6679.

111. Pfeffer, L. M. 2011. The role of nuclear factor kappaB in the interferon response. J Interferon Cytokine Res 31: 553–559.

112. Kim, D. W., L. Gazourian, S. A. Quadri, R. Romieu-Mourez, D. H. Sherr, and G. E. Sonenshein. 2000. The RelA NF-kB subunit and the aryl hydrocarbon receptor (AhR) cooperate to transactivate the c-myc promoter in mammary cells *Equal contributions. Oncogene 19: 5498–5506.

113. Vogel, C. F., E. Sciullo, W. Li, P. Wong, G. Lazennec, and F. Matsumura. 2007. RelB, a New Partner of Aryl Hydrocarbon Receptor-Mediated Transcription. Mol Endocrinol 21: 2941–2955.

114. Inozume, T., K. Hanada, Q. J. Wang, M. Ahmadzadeh, J. R. Wunderlich, S. A. Rosenberg, and J. C. Yang. 2010. Selection of CD8+PD-1+ lymphocytes in fresh human melanomas enriches for tumor-reactive T cells. J Immunother 33: 956–964.

115. Simon, S., V. Vignard, L. Florenceau, B. Dreno, A. Khammari, F. Lang, and N. Labarriere. 2016. PD-1 expression conditions T cell avidity within an antigen-specific repertoire. Oncoimmunology 5: e1104448.

116. Gros, A., M. R. Parkhurst, E. Tran, A. Pasetto, P. F. Robbins, S. Ilyas, T. D. Prickett, J. J. Gartner, J. S. Crystal, I. M. Roberts, K. Trebska-McGowan, J. R. Wunderlich, J. C. Yang, and S. A. Rosenberg. 2016. Prospective identification of neoantigen-specific lymphocytes in the peripheral blood of melanoma patients. Nat Med 22: 433–438.

