## Supplemental Figures and Legends for "The Aryl Hydrocarbon Receptor Controls IFNγ-Induced Immune Checkpoints PD-L1 and IDO via the JAK/STAT Pathway in Lung Adenocarcinoma"

### Supplemental Figure 1

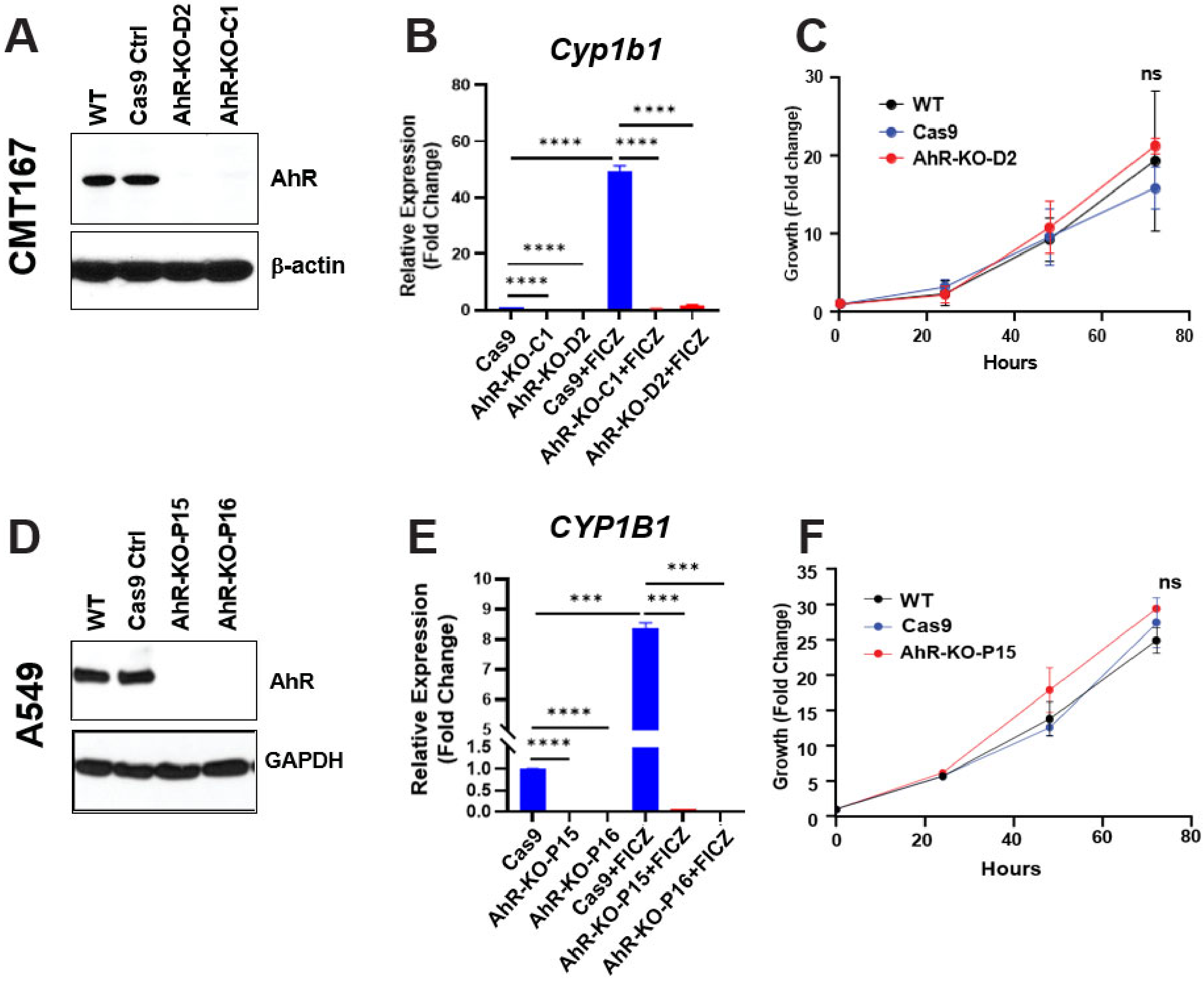


**Supplemental Figure 1. Verification AhR knockout in CMT167 and A549 cells.** CRISPR/Cas9 gene editing was used to generate two CMT167 AhR knockout clones, D2 and C1, and two A549 AhR knockout clones, P16 and P15. **A,D)** Extracted protein was evaluated for AhR expression by western immunoblotting. Data are from one of three representative experiments for each cell line. **B,E)** $CMT167^{Cas9}$, $CMT^{AhR\text{-}KO}$ (clone C1), $A549^{Ca9}$, and $A549^{AhR\text{-}KO}$ (clone P15) cells were treated with vehicle (0.1% DMSO) or 0.5 µM FICZ as an AhR activator. Twenty-four hours later extracted RNA was assayed for expression of prototypical AhR target gene *CYP1B1* by RT-qPCR. Data are from one of three independent representative experiments, each in triplicate. ***p<0.001, ****p<0.0001 (Student's t test, equal variance). **C,F)** Growth of $CMT167^{WT}$, $CMT167^{Cas9}$, $CMT^{AhR\text{-}KO}$ (clone C1), $A549^{WT}$, $A549^{Ca9}$, and $A549^{AhR\text{-}KO}$ (clone P15) cells was tracked over 72hr. Data are presented as fold increase + SE from two independent experiments, each in duplicate. ns=not significant (Student's t-test, equal variance).

**Supplemental Figure 2**

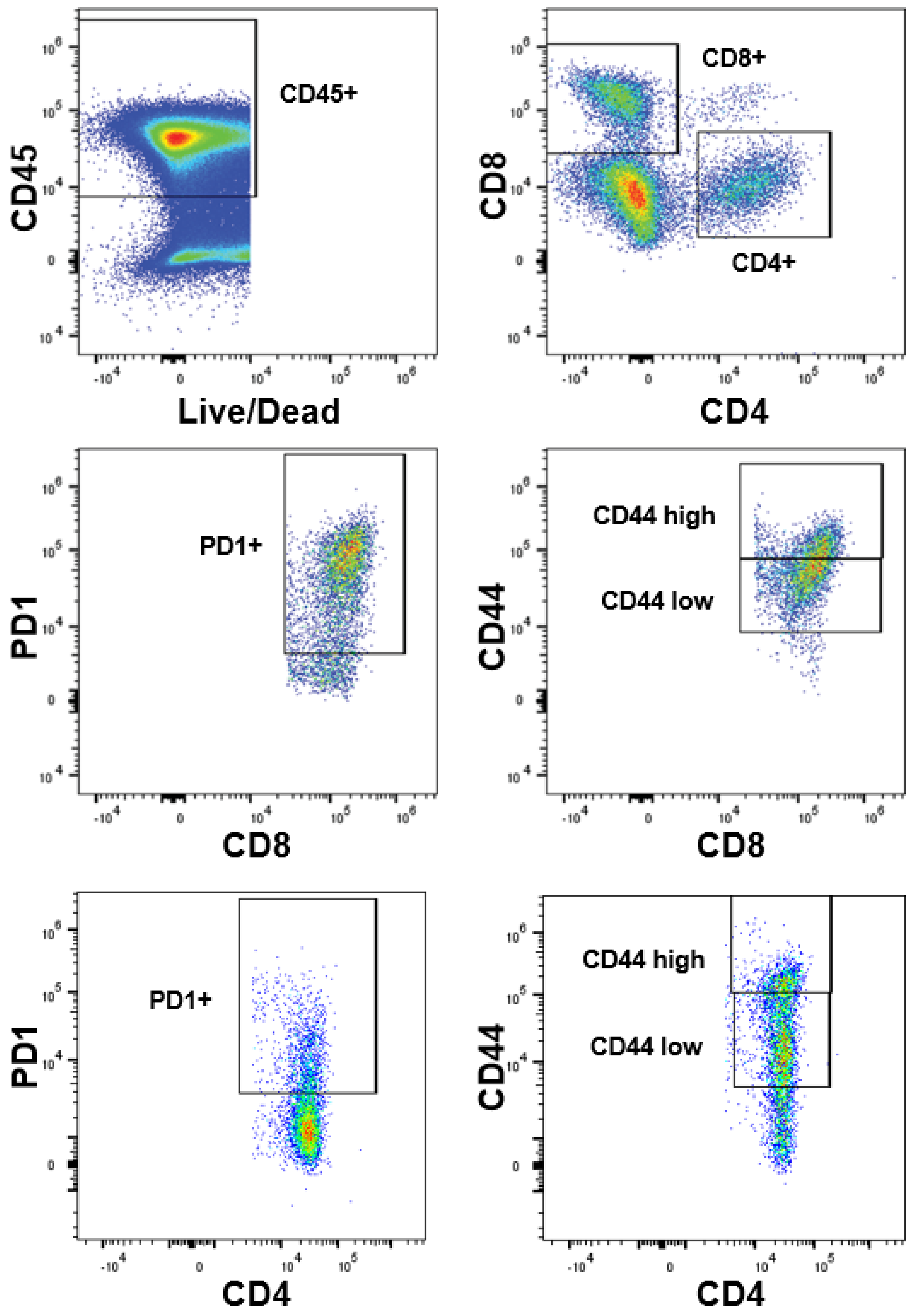


**Supplemental Figure 2. Flow cytometry gating strategy.** TILs from CMT167$^{WT}$ tumors were first gated for singlets. Lower bounds of CD4$^{+}$, CD8$^{+}$, PD-1$^{+}$, and CD44$^{+}$ cells were set with fluorescence minus one controls (FMOs). High CD44 expression was defined as above $10^5$ and low expression as below $10^5$ relative fluorescence intensity.

**Supplemental Figure 3**

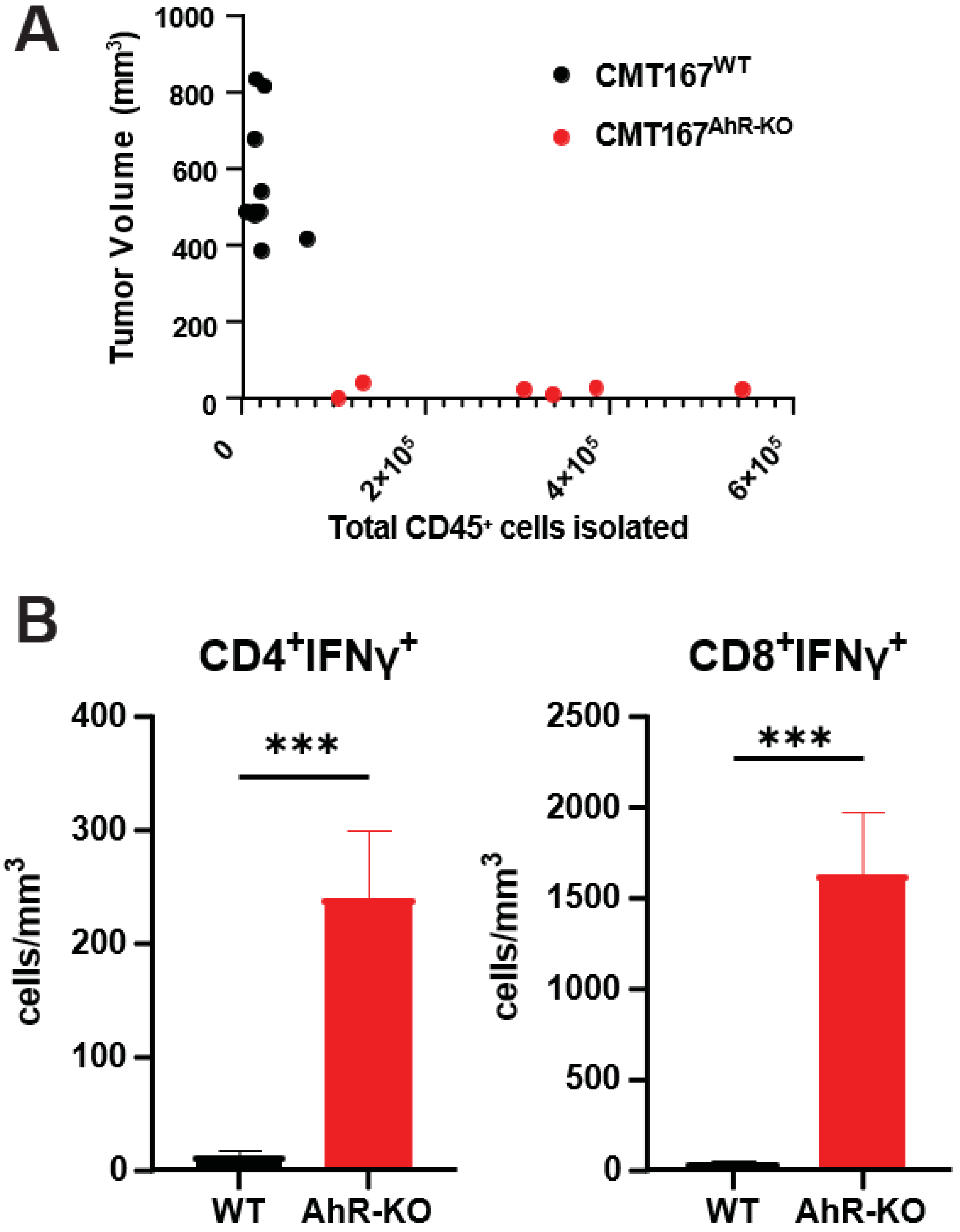


**Supplemental Figure 3. Tumor size inversely correlates with total numbers of tumor infiltrating CD45$^+$ cells. A)** The volume of CMT167$^{WT}$ and CMT167$^{AhR\text{-}KO}$ tumors (when present) were measured five weeks after transplantation. Tumors were digested, and recovered cells counted. The percent CD45$^+$ cells was determined by flow cytometry. That number was divided by the volume of the respective contributing tumors to obtain CD45$^+$ cell density (cells/mm$^3$). Data were generated from two independent experiments with totals of eight wildtype and six AhR-KO tumors. **B)** Comparison of the number of CD4$^+$IFNγ$^+$ cells/mm$^3$ (left) and CD8$^+$IFNγ$^+$ cells/mm$^3$ (right) from 12 CMT167$^{WT}$ (black) and 11 CMT167$^{AhR\text{-}KO}$ (red) tumors in two independent experiments *$p<0.05$ (Student's t-test).
